# Parkinson Disease-Linked Parkin Mediates Redox Reactions That Lower Oxidative Stress In Mammalian Brain

**DOI:** 10.1101/2020.04.26.062380

**Authors:** Daniel N. El Kodsi, Jacqueline M. Tokarew, Rajib Sengupta, Nathalie A. Lengacher, Andy C. Ng, Heather Boston, Qiubo Jiang, Carina Palmberg, Chantal Pileggi, Bojan Shutinoski, Juan Li, Angela P. Nguyen, Travis K. Fehr, Doo Soon Im, Steve Callaghan, David S. Park, Matthew J. LaVoie, Jennifer A. Chan, Masashi Takanashi, Nobutaka Hattori, Rajiv R. Ratan, Luigi Zecca, Lawrence Puente, Gary S. Shaw, Mary-Ellen Harper, Arne Holmgren, Julianna J. Tomlinson, Michael G. Schlossmacher

## Abstract

We recently hypothesized that parkin plays a role in redox homeostasis and provided evidence that it directly reduces hydrogen peroxide (H_2_O_2_) *in vitro*. Here, we examined this anti-oxidant activity *in vivo*. Informed by findings in human brain, we demonstrate that elevated oxidative stress promotes parkin insolubility in mice. In normal mouse brain parkin was partially oxidized, *e.g*., at cysteines 195 and 252, which was augmented by oxidative stress. Although under basal conditions H_2_O_2_ levels were unchanged in adult *prkn*^*-/-*^ brain, a parkin-dependent reduction of cytosolic H_2_O_2_ was observed when mitochondria were impaired, either due to neurotoxicant exposure (MPTP) or *Sod2* haploinsufficiency. In accordance, markers of oxidative stress, *e.g*., protein carbonylation and nitrotyrosination, were elevated in the cytosol but not in mitochondria from *prkn*^*-/-*^ mice. Nevertheless, this rise in oxidative stress led to changes in mitochondrial enzyme activities and the metabolism of glutathione in cells and mammalian brain. In parkin’s absence reduced glutathione concentrations were increased including in human cortex. This compensation was not due to new glutathione synthesis but attributed to elevated oxidized glutathione (GSSG)-reductase activity. Moreover, we discovered that parkin also recycled GSSG to its reduced form. With this reaction, parkin became S-glutathionylated, *e.g*., at cysteines 59 and human-specific 95. This oxidative modification was reversed by glutaredoxin. Our results demonstrate that cytosolic parkin mediates anti-oxidant reactions including H_2_O_2_ reduction and glutathione regeneration. These reducing activities lead to a range of oxidative modifications in parkin itself. In parkin-deficient brain oxidative stress rises despite changes to maintain redox balance.

## INTRODUCTION

Parkinson disease (PD) is a progressive, heterogeneous disorder of the human brain that remains incurable. Young-onset, autosomal-recessive PD (ARPD) due to deficiency in parkin (encoded by *PRKN*) is restricted to the degeneration of dopamine producing neurons in the *S. nigra* and *L. coeruleus*^1^. In the ageing human brain, these neurons are susceptible to degeneration because of unique features. These include: extensive arborization; a high number of axonal mitochondria; the presence of metals in redox-reactive forms^2^; ongoing generation of toxic dopamine metabolites in the cytosol; a greater need to buffer Ca^2+^ ions; and collectively, a greater degree of oxidative stress^3-5^.

Parkin is a RING-carrying protein with ubiquitin ligase activity that is involved in the regulation of mitophagy and immune-related functions^6,7,8^. The role for each of these functions in relation to ARPD development remains unknown. We recently described that parkin itself is highly oxidized in human brain, and discovered that it acts as a redox molecule to decrease dopamine metabolism-associated stress^9,10^.

Oxidative stress and mitochondrial damage have been implicated in the pathogenesis of several brain disorders including PD^11^. Mitochondrial dysfunction, as induced by the neurotoxicants 1-methyl-4-phenyl-1,2,3,6-tetrahydropyridine (MPTP) and rotenone, augments oxidative stress^12^. The integrity of the cellular thiol pool, a network formed by glutathione and the cysteine proteome, is essential in maintaining redox homeostasis and thus to the survival of long-lived neurons. The reduced form of glutathione (GSH) plays a critical role as an anti-oxidant, as co-factor of enzymes, and as a reservoir of cellular thiols. Accordingly, a decline in GSH has been implicated in many human disorders including neurodegenerative diseases^13^.

Genomic deficiency in parkin in mice does not induce loss of dopamine neurons, and *prkn*^*-/-*^ animals do not develop parkinsonism. However, unbiased studies have revealed proteomic changes in these animals^14,15^. There, we noted that a majority of dysregulated proteins in the brain, which showed altered isoelectric points on 2D-gels, had been previously identified as enzymes (55/92; 59.8%). Many of the proteins were also known as redox-sensitive proteins (72/92; 78.2%), *e*.*g*., peroxiredoxins, glyoxalase and aconitase-2^14,15^. These findings suggested to us that posttranslational modifications, rather than changes in protein abundance, could have explained their dysregulation.

When taken together with previous reports of parkin’s sensitivity to pro-oxidants^9,16^, we hypothesized that: one, thiol-rich parkin is altered in response to cellular redox changes; two, parkin itself plays a role in lowering oxidative stress *in vivo*, thus influencing redox-state and redox-sensitive enzymes; and three, the loss of parkin’s anti-oxidant effects contributes to mitochondrial changes. We therefore characterized parkin’s redox activity *in vitro*, explored the downstream oxidative changes in cells and mouse models with low or absent parkin expression, and investigated parkin’s effects on the thiol network. This work was informed by parallel efforts, which focused on parkin’s processing in human midbrain and its role in dopamine metabolism^10^.

## RESULTS

### Mitochondrial and cytosolic stress conditions lead to oxidation of cellular parkin

Using cell-based paradigms, we first analyzed parkin’s response to cytosolic and mitochondrial stressors. Chinese hamster ovary cells that stably express wild-type (WT) human *PRKN* cDNA (CHO-parkin) and control cells, which in our hands showed little endogenous parkin by routine Western blotting, were exposed to different levels of hydrogen peroxide (H_2_O_2_, 0.2 or 2mM) or carbonyl-cyanide-m-chlorophenyl-hydrazine (CCCP, 10μM). Under these stress conditions, which did not cause cell death, we observed the time-dependent loss of parkin’s monomer (∼52.5 kDa) and an increase in high molecular weight (HMW) species (**Fig. 1a**)^16^. These biochemical changes were seen under non-reducing electrophoresis conditions and thus interpreted as involving parkin thiol-based oxidation. Intriguingly, this oxidation was specific to parkin as endogenous, redox-sensitive and ARPD-linked Dj-1^17^ did not undergo such changes (**Fig. 1a**).

**Figure 1:**
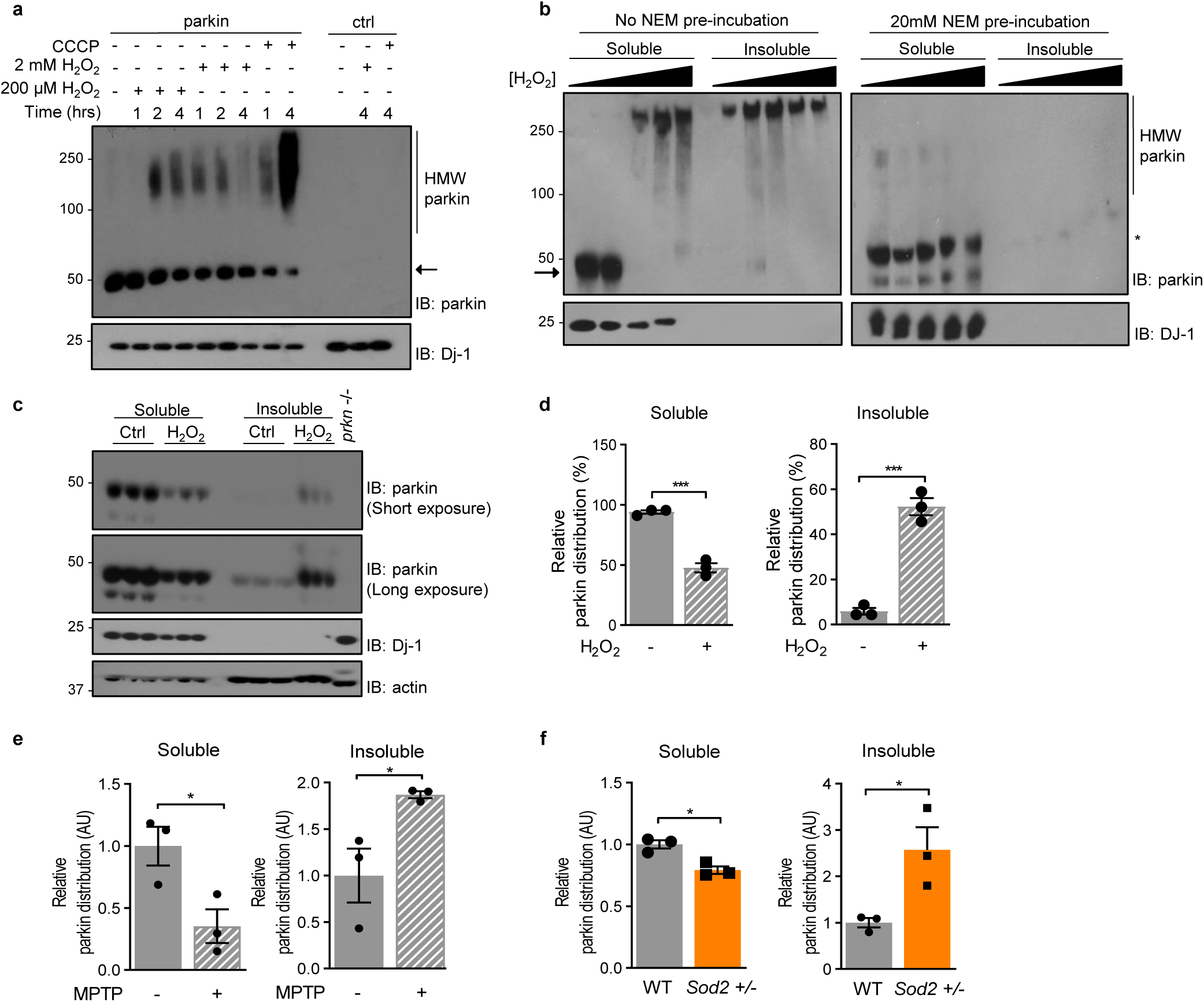
Mitochondrial and cytosolic stressors lead to oxidation of cellular parkin promoting its insolubility. Representative Western blots of parkin and Dj-1 distribution in, **a**, CHO-parkin and control cells under oxidizing conditions (200μM and 2mM H_2_O_2_; and 10μM CCCP), **b**, soluble and insoluble fractions from HEK293-parkin and control cells incubated with increasing concentrations of H_2_O_2_ (2μM to 2M) with or without 20mM n-ethylmaleimide (NEM) pre-incubation, and **c**, soluble and insoluble fraction from 2-4 mths-old wild-type C57Bl/6 mouse brains incubated with saline or 1% H_2_O_2_. (←) Monomeric parkin (*) Alkylated-parkin. Quantification of parkin’s signal distribution in brain homogenates from, **d**, 2-4 mths-old wild-type C57Bl/6 mouse brains incubated with saline or 1% H_2_O_2_, **e**, 2-4 mths-old wild-type C57Bl/6 mice treated with a 40mg/kg intraperitoneal injection of MPTP toxin or saline. **f**, 6-8 mths-old C57Bl/6 wild-type (WT) and *Sod2*^+/-^ mouse brains (soluble and insoluble fractions are shown). A Student t-test was used for statistical analysis (* = < 0.05, *** = <0.001).

In oxidative stress-exposed, human embryonic kidney (HEK293) cells, which transiently over-expressed *PRKN*, we also observed HMW smear formation. Parkin became progressively gel excluded (aggregated) and insoluble following exposure to increasing concentrations of H_2_O_2_ (**Fig. 1b**). Since these HMW forms of parkin were reversed with dithiothreitol (DTT; **Extended Data Fig. 1a, c**), we postulated that these modifications occurred due to increasing oxidation of parkin’s cysteines^16^. We tested this using N-ethylmaleimide (NEM) and iodoacetamide (IAA), which irreversibly alkylate free thiols. There, we found that pre-incubation of cells with NEM or IAA blocked HMW smear formation of parkin and preserved its solubility during H_2_O_2_ stress (**Fig. 1b; Extended Data Fig. 1b**). Complementing these findings, we mapped several cysteines in recombinant (r-) parkin, which became oxidized when the protein was subjected to rising H_2_O_2_ levels *in vitro*, thereby promoting its aggregation and insolubility^10^.

Intriguingly, exposure to DTT also lowered parkin solubility in cells (**Extended Data Fig. 1c**), which suggested that parkin’s structure is altered under excessive oxidizing and excessive reducing conditions. These results demonstrated that human parkin undergoes thiol chemistry-based changes in cells during redox stress.

### Excessive ROS exposure in mice promotes parkin insolubility

Under stress-free conditions, parkin resides in the cytosol of cells (**Fig. 1a, b**) and rodent brain^18,19^. In contrast, in human control brain parkin is largely insoluble after the 4^th^ decade of life^10,19^. To better understand the link between parkin’s oxidation and insolubility, we sought to validate these findings in mice in three different ways. First, freshly dissected brains from WT mice were homogenized either in the presence or absence of H_2_O_2_ to generate high amounts of reactive oxidative species (ROS). Lysates were then fractionated and separated into soluble *vs*. insoluble fractions; in the latter, proteins were extracted using 2-10% SDS. Exposure to H_2_O_2_ *in situ* led to a significant loss in solubility of mouse parkin and recovery in the insoluble fraction (P<0.001), as monitored by reducing SDS/PAGE. No such shift was seen for murine Dj-1 (**Fig. 1c, d**).

Secondly, we examined parkin’s response to oxidative stress *in vivo*. We injected a single dose of MPTP into the peritoneum of adult mice at a dose (40mg/kg) that induces oxidative stress without neuronal death^20^. After one hour, brains were collected and serially fractionated. The brief exposure to MPTP led to a decline of >50% in soluble parkin when compared to saline-treated controls. As expected, parkin’s detection rose in the insoluble fraction (P<0.05; **Fig. 1e; Extended Data Fig. 1d, e**).

In a third approach, we tested whether a genetically induced, mitochondrial defect altered parkin solubility. We chose *Sod2*^*+/-*^ haploinsufficient mice with lower expression of mitochondrial superoxide dismutase-2 (aka MnSOD). Reduced MnSOD2 activity (**Extended Data Fig. 1h, i**) results in systemically elevated oxidative stress^21^. As expected, parkin solubility had shifted in adult *Sod2*^*+/-*^ brain (P<0.05; **Fig. 1f**; **Extended Data Fig. 1j, k**). We concluded that toxicologically and genetically induced, oxidative stress led to parkin insolubility in mammalian brain. We postulated that this change corresponded with fewer free thiols, due to increasing oxidation, within parkin itself.

### Exposure to MPTP increases parkin oxidation in the brain

To test this, we employed liquid chromatography and mass spectrometry (LC-MS/MS). Brains from mice injected with MPTP or saline (as above) were homogenized in the presence of IAA to alkylate free cysteines and prevent thiol oxidation during processing. Parkin-enriched isolates were separated by SDS/PAGE (**Extended Data Fig. 1f**). Monomeric, alkylated parkin was gel-excised, trypsin digested and subjected to LC-MS/MS. We recorded a <20% decline in the relative amount of unmodified cysteines in parkin peptides from MPTP-treated mice compared to those from saline-injected controls (**Extended Data Table 1**). Furthermore, we found that several cysteines had been oxidized under normal conditions, *e*.*g*., to dehydroalanine, sulfinic acid and sulfonic acid (**Extended Data Table 1**). Intriguingly, residues C195 and C252 were frequently identified as irreversibly oxidized in mouse brain (**Extended Data Table 1; Extended Data Fig. 1g**). In parallel studies of human parkin, we found that the oxidation of orthologous cysteines, *e*.*g*., C253, contributed to its progressive insolubility^10^.

### Parkin reduces hydrogen peroxide to water in a thiol-dependent manner

We recently provided evidence that WT r-parkin reduced H_2_O_2_ to water *in vitro* by reciprocal oxidation of its thiol groups^10^; ARPD-linked point mutants did so less well. Here, we further explored the biochemical mechanisms underlying this activity. Human ‘r-parkin was nearly 5-fold more effective in lowering H_2_O_2_ levels than GSH at equimolar concentrations, but was less potent than catalase (**Extended Data Fig. 2a)**. This suggested that parkin shared a thiol-based activity with GSH. We confirmed this by demonstrating that r-parkin’s ability to reduce H_2_O_2_ was abrogated when its thiols were either oxidized prior to the reaction or irreversibly blocked by NEM (**Extended Data Fig. 2b, c**). Parkin did not exhibit peroxidase-type anti-oxidant activity (**Supp. Fig. 2D**).

Two other PD-linked proteins, *e*.*g*., α-synuclein and DJ-1 (with 0 and 3 cysteines, respectively; **Extended Data Fig. 2e**), had no detectable H_2_O_2_-lowering activity (**Fig. 2a**). Bovine serum albumin with the same number of cysteines as human parkin (n, 35) and two RING-carrying E3 ubiquitin ligases, *e*.*g*., RNF43 and HOIP (**Extended Data Fig. 2e, f**), also failed to lower H_2_O_2_ (**Fig. 2a**). The latter results suggested that parkin’s structure played a role in its activity, which was not solely due to the presence of its four RING domains, each of which chelates two Zn^2+^ ions (**Extended Data Fig. 1g**)^22^. Nevertheless, competitive chelation of Zn^2+^ ions by rising EDTA concentrations prior to the reaction diminished r-parkin’s antioxidant activity. This was partially, but not completely, reversed when excess Zn^2+^ was added back to the reaction mix (**Extended Data Fig. 2g**). Of note, EDTA and NEM alone had no effect on the assay (**Extended Data Fig. 2h**). We concluded from these findings that parkin’s H_2_O_2_-reducing activity was: not enzymatic, required integrity of its tertiary structure, depended on free thiols, and resembled GSH’s mechanism of action.

**Figure 2:**
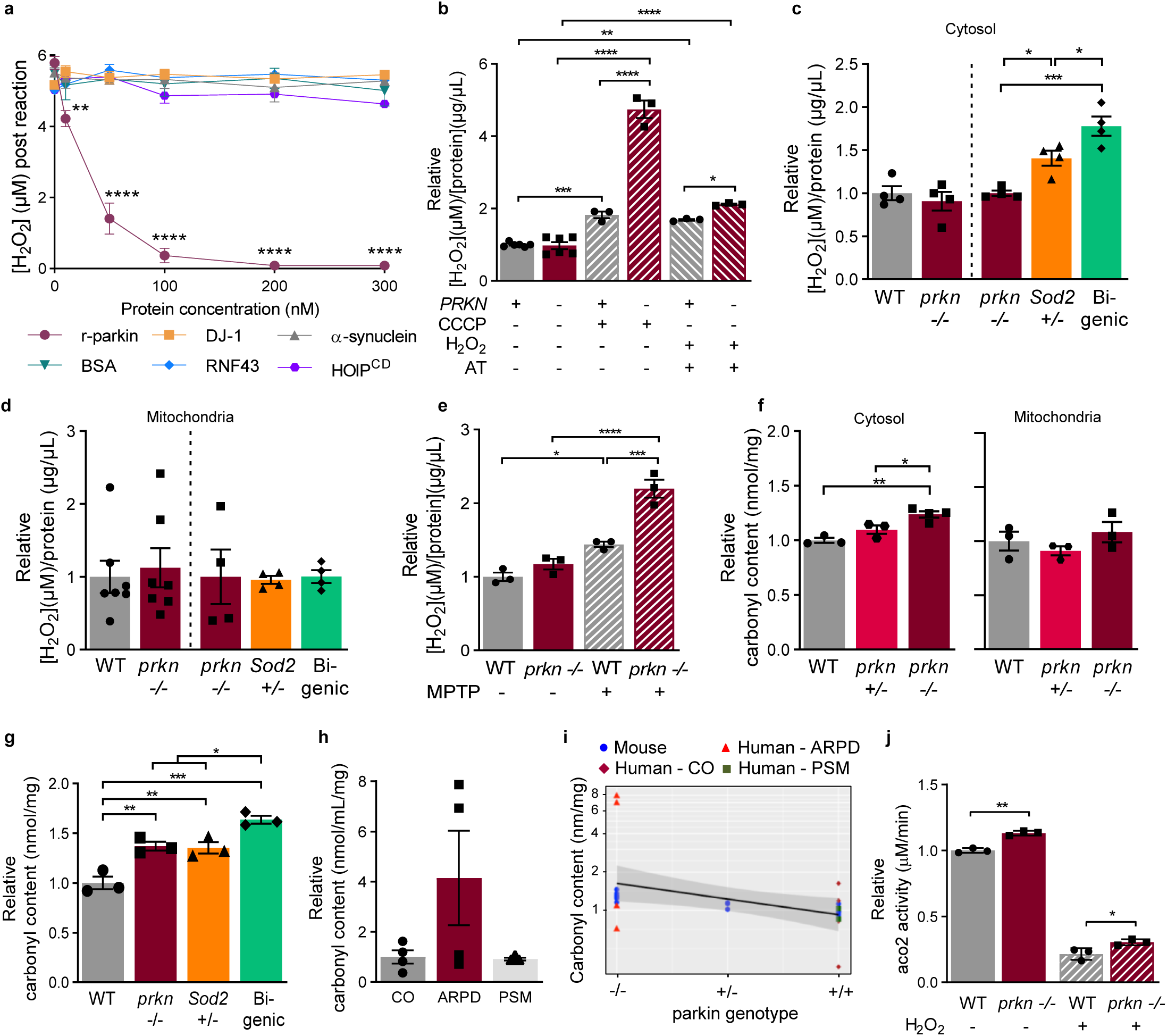
Parkin cysteines reduce hydrogen peroxide to lower oxidative stress. **a**, Parkin’s reducing capacity compared to DJ-1, α-synuclein, bovine serum albumin (BSA), and two RING-carrying ubiquitin ligases (RNF43 and HOIP^cd^, (^cd^ - catalytic domain)). **b**, Endogenous levels of H_2_O_2_ in HEK293-parkin and control cells under control *vs*. oxidizing conditions [10μM CCCP for 1hr; 0.5M AT (to inhibit catalase) + 2mM H_2_O_2_ for 30mins]. Endogenous levels of H_2_O_2_ in **c**, cytosol-enriched and in **d**, mitochondria-rich brain homogenates of 6-8 mths-old WT and *prkn*^-/-^ and, 2-4 mths-old *prkn*^-/-^, *Sod2*^+/-^ and bi-genic (*prkn*^-/-^//*Sod2*^+/-^) C57Bl/6 mice, and in **e**, 6-8 mths-old WT or *prkn*^-/-^ mouse brains treated *in vivo* with either saline or MPTP toxin. Protein carbonyl content in **f**, cytosol-enriched and mitochondria-rich brain homogenates from 8 mths-old WT, *prkn*^+/-^, and *prkn*^-/-^ mouse brains, **g**, brain lysates of 6 mths old WT, *prkn*^-/-^, *Sod2*^+/-^ and bigenic mice, and **h**, aged-matched cortex specimens from controls, *PRKN*-deficient ARPD patients and subjects with other forms of parkinsonism (PSM). **i**, Carbonyl content (nmol/mL/mg) in mammalian brains of different *PRKN* genotypes. Linear regression line (blue) with standard error (gray shade) signals the overall trend. **j**, Aconitase-2 enzyme activity in mitochondria isolated from 12 mths-old WT and *prkn*^-/-^ mouse brains. 1-way or 2-way ANOVA was used for statistical analysis (* = < 0.05; ** = < 0.01; *** = < 0.001; and ****= <0.0001).

### *PRKN* expression lowers hydrogen peroxide concentrations in the cytosol

We next asked whether parkin contributed to redox balance in cells and *in vivo*. Under basal conditions, HEK293-parkin cell lysates showed a trend toward lower levels of H_2_O_2_ when compared to controls; the difference became significant when we monitored intact cells (**Fig. 2b**; **Extended Data Fig. 2i**). When HEK293 cultures were exposed to rising oxidative stress, *e*.*g*., after CCCP treatment or excessive levels of H_2_O_2_ in the presence of aminotriazole (AT, a catalase inhibitor; **Extended Data Fig. 2a**), parkin-expressing cell lysates showed a significantly lower level of ROS (P<0.0001; **Fig. 2b**).

We next tested parkin’s effects on H_2_O_2_ concentrations in mouse brain using a validated AmplexRed assay (**Extended Data Fig. 2j**). We found that basal ROS concentrations did not differ between adult *prkn*^*-/-*^ and WT mice, neither in the cytosol nor in isolated mitochondria (**Fig. 2c, d**; **Extended Data Fig. 3b**). In contrast, exposure to MPTP caused a significant elevation in endogenous ROS levels in parkin-deficient mice (P<0.001; **Fig. 2e**). Under basal conditions H_2_O_2_ levels were also higher in the cytosol of *Sod2*^*+/-*^ mutant and bigenic animals (*prkn*^*-/-*^*//Sod2*^*+/-*^), which we had created to further augment metabolic stress (P<0.05; **Fig. 2c**; **Extended Data Fig. 1h, i**). Of note, at this age *Sod2*^*+/-*^ and bigenic mice did not show detectable cell loss (not shown). Unexpectedly, ROS levels remained the same in isolated brain mitochondria from *Sod2*^*+/-*^ mice (**Fig. 2d**), which suggested rapid shuttling of superoxide from their mitochondria into the cytosol^23^.

### *Prkn* gene expression lowers oxidative stress in mammalian brain

We next examined the degree of protein carbonylation, a marker of chronically elevated H_2_O_2_, in the same mouse lines. Carbonyl content was increased in the cytosol of *prkn-* mutant brain, even under basal conditions, consistent with a previous report by Palacino et al., who had used a different mouse model^14^, but remained normal in isolated mitochondria. There, we also observed a *prkn-*null allele dosage effect (**Fig. 2f**). As expected, carbonyl content was further increased in the bigenic mice when compared to WT, *prkn*^-/-^ and *Sod2*^+/-^ littermates (P<0.001 and P<0.05 respectively; **Fig. 2g**). A second marker of oxidative stress-induced protein modification, *i*.*e*., nitrotyrosination, was also increased in the cytosol of *prkn*^*-/-*^ mice including in the heart (**Extended Data Fig. 3c, d**).

We next examined protein carbonyl content in patients with *PRKN*-linked ARPD and control subjects (each, n=4). These had been matched for age, *post mortem* interval, ethnicity and hospital morgue (**Extended Data Fig. 3a**)^24^. In the absence of human parkin, protein carbonyl levels showed a trend toward elevation. Cortices from three, non-*PRKN-*linked parkinsonism cases revealed the same degree of carbonyl content as age-matched controls (**Fig. 2h**). Using a linear regression model with the protein carbonyl content measured in all brains (mouse and human) as the dependent variable, we found that oxidative stress correlated with parkin deficiency; the co-efficient was −0.3754 (95% CI, −0.6611–0.0053; P=0.05; **Fig. 2i**). Of note, in these redox studies, we did not observe any sex effect. Together, these results established that WT parkin contributed to H_2_O_2_ reduction in cells and brain during basal activities and with rising oxidative stress.

### *Prkn* gene expression affects redox-dependent enzymatic activity

To link these findings back to proteomic changes downstream of parkin deficiency, we explored redox-dependent enzymatic activities. In the brains of *prkn* ^*-/-*^ mice, we found significantly increased activity of aconitase-2 (Aco2), a NADP^+^-linked enzyme, under basal conditions when compared to littermates (P<0.001; **Fig. 2j**). Following exposure of isolated brain mitochondria to H_2_O_2_, Aco2 and creatine kinase (mtCK) activities were decreased in both WT and *prkn*-null brains, as expected; however, their activities remained measurably higher in *prkn*^*-/-*^ mitochondria (P<0.05 and P<0.01, respectively; **Fig. 2j**; **Extended Data Fig. 3e**). Importantly, the levels of detectable murine and human Aco2 and mtCK proteins were not altered in these brains, neither in whole lysates nor in isolated mitochondria, when comparing WT *vs*. mutant *PRKN* genotypes (**Extended Data Fig. 3b, f-h**). Although increased cytosolic ROS production in parkin-deficient brain could have led to the proteomic changes detected in these animals, this would not have fully explained altered enzyme activities by Aco2 and mtCK under redox stress. We suspected that other redox effects in *prkn*^*-/-*^ mice, such as changes in the thiol network, contributed to these results.

### Parkin contributes to the thiol network during oxidative stress

To test whether parkin expression contributed to the thiol network, we first compared survival rates of CHO-parkin cells following exposure to H_2_O_2_ after depletion of GSH with buthionine sulfoximine (BSO). Treatment with either H_2_O_2_ (1h) or BSO alone (48hs) did not cause cell death (**Extended Data Fig. 2k**). Under combined stress conditions, *PRKN* expression was protective: it led to significantly less cell death (P<0.05; **Fig. 3a**) and normalized ROS levels (P<0.001; **Fig. 3b**), resulting in reciprocal parkin oxidation (Fig. 1A). Treatment with excess n-acetylcysteine, an abundant source of free thiols, resulted in similarly protective effects as did *PRKN* expression (**Fig. 3a, b**). Based on these results, we conjectured that parkin supplements the cellular thiol network through its own free thiols, which could neutralize ROS.

**Figure 3:**
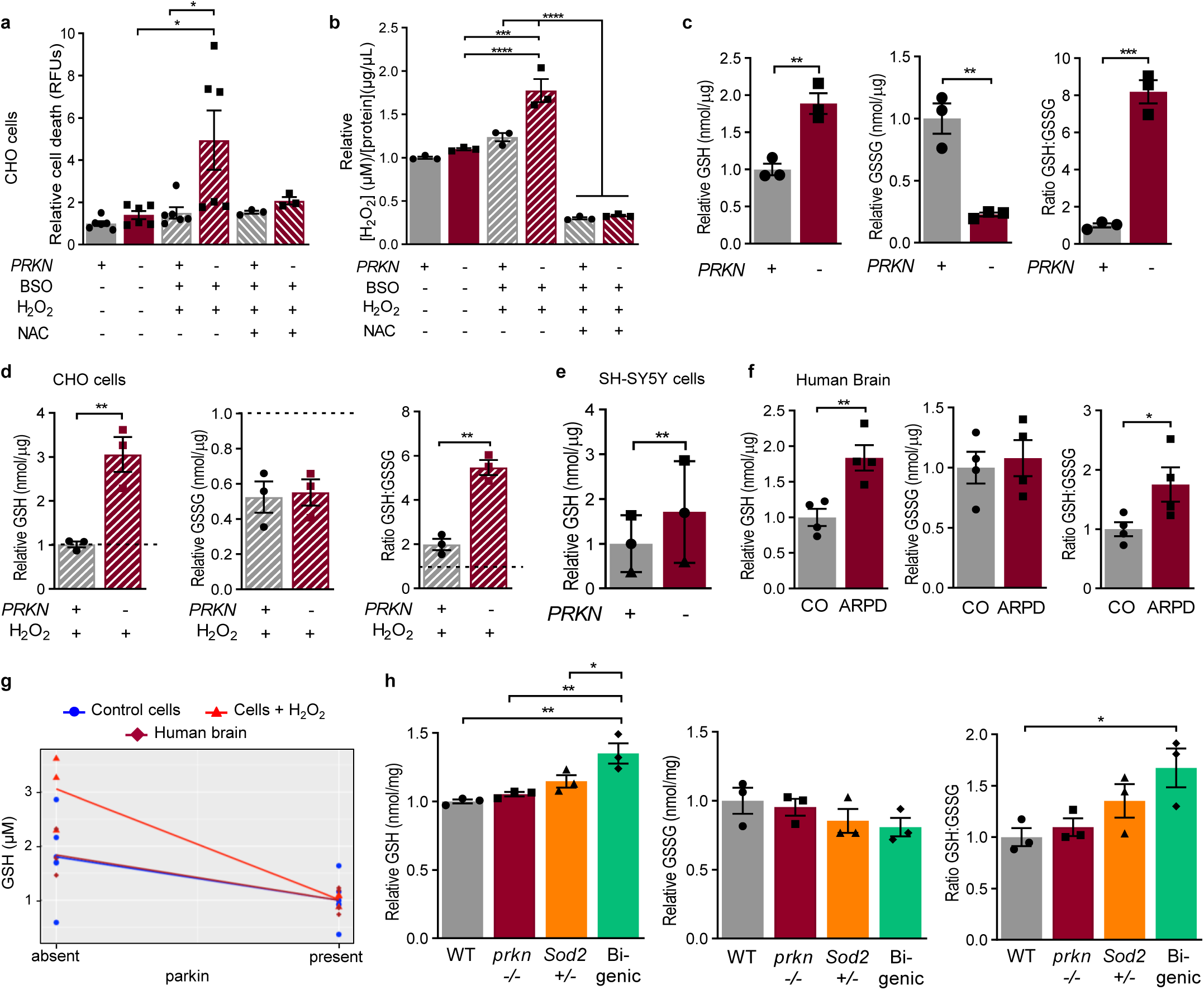
*PRKN* expression alters glutathione metabolism in cells and human brain. Cellular toxicity **a**, and endogenous H_2_O_2_ levels **b**, in CHO-parkin and control cells under control or oxidizing conditions (2 mM H_2_O_2_ + 2mM buthionine sulfoximine (BSO)), with or without supplementation by 20mM n-acetylcysteine (NAC). Reduced GSH, oxidized GSSG, and GSH:GSSG ratio in CHO-parkin or control cells under **c**, control conditions, and **d**, post H_2_O_2_ stress (HPLC method). **e**, GSH levels in SHSY5Y cells under control conditions (monochlorobimane (MCB) method). Paired data are represented by a distinct symbol. GSH, GSSG, GSH:GSSG ratio in **f**, control and *PRKN* deficient human brain cortices, and **h**, brains of 6 mths-old WT, *prkn* -/-, *sod2* +/- and bi-genic mice (HPLC method). **g**, GSH (μM) levels in several paradigms of *PRKN* genotype comparisons, as indicated. Linear regression lines represented the overall trend of control cells (blue), cells + H_2_O_2_ (red), and human brain (darker red). A Student t-test or 1-way unpaired or paired ANOVA were used for statistical analysis (* = < 0.05; ** = < 0.01; *** = < 0.001; and ****= <0.0001).

### *PRKN* gene expression alters glutathione redox state in cells and human brain

We examined this by directly measuring levels of GSH (reduced) and GSSG (oxidized) by HPLC. There, CHO-control cells had relatively high concentrations of GSH and low amounts of GSSG, with an elevated ratio of GSH:GSSG (**Fig. 3c**). Increased expression of WT parkin significantly reversed these redox indices (P<0.01 and P<0.001), but no change was found in the total concentration of GSH and GSSG (**Extended Data Fig. 4a**). Consistent with reports of CHO cells displaying high stress tolerance^25^, exogenously added H_2_O_2_ did not substantially change the redox indices between CHO-control and CHO-parkin cells with the exception of now equalized GSSG levels (**Fig. 3d**). Parkin-dependent changes in GSH levels and GSH:GSSG ratios were also observed in transfected HEK293 cells (not shown) and human, dopamine SH-SY5Y cells (**Fig. 3e**).

The interplay between *PRKN* expression and GSH redox state was further validated in human brain using cortex specimens from patients with *PRKN*-linked ARPD and healthy controls (as used above). In the absence of detectable parkin, reduced GSH levels as well as the ratio of GSH:GSSG were significantly increased (P<0.01 and P<0.05 respectively). No differences were seen for GSSG levels (**Fig. 3f**) or the total concentration of GSH and GSSG (**Extended Data Fig. 4b**). Intriguingly, the redox changes recorded in human brain closely mirrored those in CHO cells exposed to oxidative stress (**Fig. 3d**). We concluded that a reciprocal relationship existed between *PRKN* expression and GSH levels *in vivo*. When combining these results and using a linear regression model with GSH levels as the dependent variable, we found that lower expression of parkin was associated with higher GSH levels; the co-efficient was −0.6889 (95% CI, −0.849 to −0.411; P<0.0001; **Fig. 3g**).

### *Prkn* deletion alters glutathione metabolism in mice with a mitochondrial defect

Two groups previously reported elevated GSH levels in *prkn* ^*-/-*^ mice and their glia^26,27^. When examining the same colony generated by Itier et al. using our HPLC method, parkin-deficient brains showed similar but non-significant trends for thiol changes, *i*.*e*. higher concentrations of reduced GSH, lower levels of GSSG, and a higher GSH:GSSG ratio (**Fig. 3h**). Brains from *Sod2*^*+/-*^ haploinsufficient animals showed identical results. When endogenous oxidative stress was further elevated in bigenic (*prkn* ^*-/-*^*//Sod2*^*+/-*^) animals, the same changes in redox indices were detected (P<0.05; **Fig. 3h**), as measured in human parkin-deficient cells and cortices (above). These results established a relation between parkin protein expression, glutathione redox state and degree of oxidative stress *in vivo*. We next studied potential mechanism(s) to explain it.

### *De novo* synthesis of glutathione is not altered in parkin-deficient brain

We first examined GSH *de novo* synthesis. When we measured the total pool of glutathione (GSH and GSSG) by HPLC in our parkin-deficient *vs*. WT parkin expression paradigms, we saw no differences for the total concentration across cell cultures, murine brains and human cortices (**Extended Data Fig. 4a, b, c**). In accordance, we found no difference in transcript numbers for the two rate-limiting enzymes in GSH synthesis, *i*.*e*., the glutamate-cysteine-ligase-catalytic (*GCLC*) and glutamate-cysteine-ligase-modifier (*GCLM*) subunits^28^, or for *Dj-1*, in these mouse lines (**Extended Data Fig. 4d, e**). Unexpectedly, in bigenic mice, where we did record a significant rise in the total glutathione (GSH + GSSG) concentration (P<0.05), we found no higher copy numbers for *GCLC* or *GCLM* mRNA (**Extended Data Fig. 4c**,**e**). Thus, *de novo* synthesis from transcription did not explain the glutathione redox changes seen in parkin’s absence.

### Glutathione recycling is altered in *prkn*-null brain

We next explored a second mechanism, namely enhanced regeneration of GSH by glutathione reductase (GR). This enzyme is activated by rising levels of ROS or its substrate (GSSG), and is inhibited by excess product levels, such as GSH or NADP^+ 29-31^. By using the Tietze method, which utilizes exogenous GR activity to indirectly measure GSH and GSSG levels, we reproduced the same significant changes, *i*.*e*., GSH elevation (P<0.01), GSSG lowering (P<0.01) and the rise of the GSH:GSSG ratio, in *prkn* ^*-/-*^ mice as found in cells and human brain (**Fig. 4a** and **Fig. 3c-f**, respectively). These findings in mouse brain matched the published results by Itier et al.; they suggested that GR activity was indeed higher in parkin’s absence. We reasoned that activation of endogenous GR was even more elevated in ARPD cortices than in *prkn*^-/-^ brain, because GSH levels were higher in the former than the latter using the GR-independent HPLC method (**Fig. 3f-h**).

**Figure 4:**
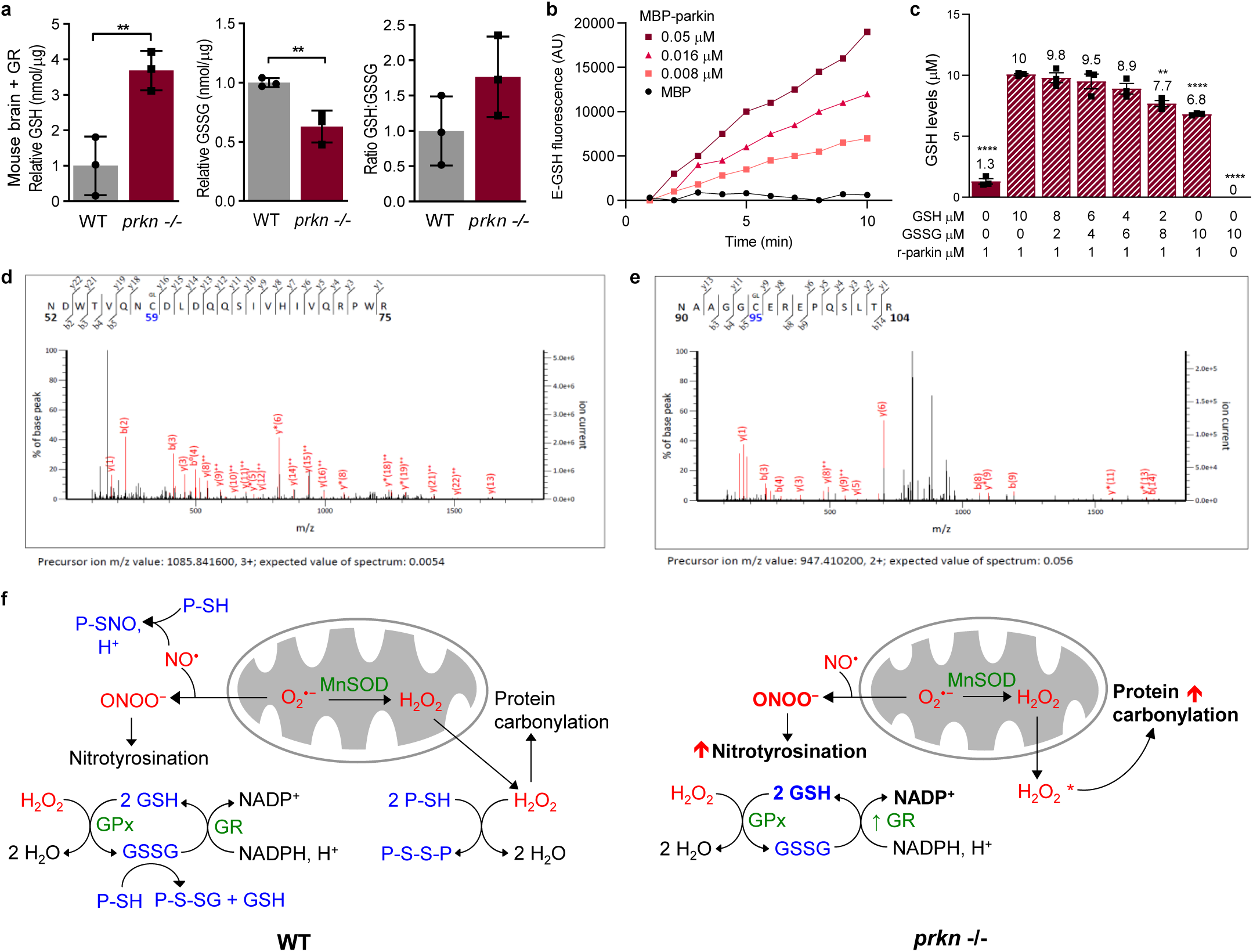
Parkin contributes to glutathione recycling independent of *de novo* synthesis. **a**, GSH, GSSG, and GSH:GSSG ratio in 11-13 mths-old WT and *prkn*^-/-^ mouse brains (GR-dependent Tietze method). **b**, E-GSH release after exposure of increasing concentrations of full-length MBP-parkin to 20μM Di-E-GSSG. **c**, GSH levels (μM) of various GSH:GSSG ratios incubated with 1μM of full-length, untagged r-parkin. A Student t-test or 1way ANOVA was used for statistical analysis (** = < 0.01; and *** = < 0.001). **d**, Human parkin peptide aa52-75 is shown, with cysteine 59 S-glutathionylated. **e**, Human parkin peptide aa90-104 is shown, with cysteine 95 S-glutathionylated. **f**, Graphical depiction of redox changes including: GSH recycling; generation of ROS levels (*i*.*e*., H_2_O_2_; superoxide) leading to carbonylation; metabolism of nitric oxide (NO) leading to nitrotyrosination; and function of select enzymes in WT *vs*. parkin-deficient (*prkn*^-/-^) mammalian brain. P, parkin; MnSOD, Mn^2+^-dependent superoxide dismutase; GPx, glutathione peroxidase; GR, glutathione reductase. * Indicates evidence from human brain^10^.

### Parkin contributes to the pool of recycled glutathione

Based on the dynamic changes observed in the redox state of cells and tissues, we also asked whether parkin had any direct effects on thiol network changes, such as via GSH recycling. We specifically asked whether parkin interacted with GSSG. We first tested this *in vitro* using recombinant proteins, both tagged and untagged, and eosin-labelled GSSG (referred to as Di-E-GSSG) (**Extended Data Figs. 2f, 4f**). We discovered that full-length, human parkin had concentration-dependent activity in reducing Di-E-GSSG to the fluorescence-emitting E-GSH (545 nm) (**Fig. 4b**). Of note, Di-E-GSSG itself is non-fluorescent due to quenching^32^. N-terminally truncated parkin comprising the IBR-RING2 domains (aa 327-465) and a C-terminal RING2 peptide (aa 425-465) also showed E-GSH-regenerating activity (**Extended Data Fig. 4f, g**; and not shown). In contrast, the control protein-tag alone (*e*.*g*., maltose-binding protein (MBP); ∼50 kDa) and incubation with E-GSH (without any Di-E-GSSG present) showed no activity in this assay (**Fig. 4b**).

Resulting from this biochemical reaction, the formation of E-S-glutathionylated parkin (referred to as parkin-SG-E) was confirmed by using MBP-tagged parkin proteins, non-reducing SDS/PAGE analyses and UV light. We found that the ∼90 kDa MBP-parkin-SG-E and the ∼60 kDa MBP-IBR-RING2-SG-E proteins were only visible under UV light after the incubation of native proteins with Di-E-GSSG (*e*.*g*., **Extended Data Fig. 4gh**, lane 2 of left panel). Furthermore, the incubation of MBP-IBR-RING2-SG-E and MBP-parkin-SG-E with either DTT or a reconstituted glutaredoxin-1 or -2 (Grx1, Grx2) system^32^ reversed the S-glutathionylation of parkin, confirming that oxidation in the form of disulphide-bond formation had occurred during the earlier reaction (**Extended Data Fig. 4h**). In accordance, parkin proteins without any -SG-E modification did not interact with Grx1, Grx2, or thioredoxin-1 (Trx1; not shown). We reasoned that parkin’s ability to directly interact with GSSG could therefore alter the redox index of the GSH:GSSG ratio.

To test this, we incubated untagged r-parkin (1μM) with eosin-free ratios of [GSH]:[GSSG] at a total concentration of 10μM per well (**Fig. 4c**). After 15 min, we measured GSH concentrations with a specific assay. Following its incubation with GSSG alone, we determined that under these conditions parkin was able to reduce up to 5 equivalents of GSSG to GSH (6.8µM GSH minus a baseline signal of 1.3µM for r-parkin alone; **Fig. 4c**). Other ratios of [GSH]:[GSSG] showed analogous changes. Therefore, for every recycled GSSG molecule one reduced GSH and one S-glutathionylated form attached to parkin had been generated. These findings demonstrated a direct biochemical effect by parkin onto its surrounding thiol network.

We next sought to map specific S-glutathionylation sites on parkin using MBP-parkin-SG preparations by mass spectrometry, which was performed under non-reducing conditions. Full-length MBP-parkin-SG-E was subjected to trypsin digestion followed by LC-MS/MS and MALDI analyses. We detected two residues that were consistently S-glutathionylated, *i*.*e*., the conserved cysteine 59 and human-specific cysteine 95; each carried an extra mass of 305.0682 (**Fig. 4d, e**), corresponding to S-linked glutathionylated oxidation. A third, but less frequently identified residue to be S-glutathionylated was cysteine 377 within parkin’s IBR domain. To date, we have not yet identified modified cysteine(s) in the C-terminal RING2 peptide, which also had shown GSSG-recycling activity. We concluded from these results that S-glutathionylation represents a *bona fide* oxidative modification of human parkin, which arises from its interaction with GSSG.

## DISCUSSION

Our study generated four novel insights into the protective effects of PD-linked parkin: One, we restaged in cell models and mouse brain the age-associated loss of parkin solubility that occurs in human brain^10,19^ by inducing its progressive oxidation using mitochondrial and cytosolic stressors (**Fig. 1; Extended Data Fig. 1**). We then monitored for specific cysteine oxidation sites in murine parkin using LC-MS/MS (**Extended Data Fig. 2f, g; Extended Data Table 1**). There, we found a relative loss of free thiols in parkin associated with the rise of oxidative stress in MPTP-treated mice. We also mapped several cysteines that had undergone irreversible oxidation, such as to sulfinic and sulfonic acid, under both basal and oxidizing conditions, *e*.*g*., C211, C195, C237, C252, and C376 (**Extended Data Table 1**). We noticed with interest –given the role of parkin’s phosphorylation by ARPD-linked PINK1 during mitophagy^33^-that S77 of murine parkin was found phosphorylated in MPTP-exposed brain (**Extended Data Table 1**).

Two, we established that *PRKN* expression contributes to cytosolic ROS neutralization and *in vivo* redox homeostasis; this, to limit oxidative stress in high energy-demanding organs, such as the brain and heart. Its function in ROS neutralization complements parkin’s role in dopamine radicals-induced stress^10^. These anti-oxidant effects are mediated by parkin’s cysteines, which are sensitive to oxidizing as well as reducing environments (**Fig. 2a; Extended Data Fig**.**2a-g**). We found that parkin contributes to redox homeostasis downstream of oxidative stress production, as seen in lower carbonyl contents of mouse and human tissues (**Fig. 2f-i**), as first described by Palacino et al.^14^. We also provided evidence that parkin’s modulation of cytosolic ROS is linked to the activities of redox-sensitive, mitochondrial enzymes (*e*.*g*., Aco2; **Fig. 2j**), which were previously found structurally altered (but mechanistically never explained) in parkin-deficient mice^15^. Linking parkin’s role in mitigating ROS concentrations in the cytosol to changes in mitochondrial activity also helps explain the observation made by Berger et al., who had postulated an ‘intrinsic, pro-mitochondrial effect by cytosolic parkin’^34^. We suspect that the structural changes in redox-sensitive proteins from *prkn*^-/-^ mice^14,15^ were mediated by the net reduction in protein-based free thiols, thereby not only conferring less protection from endogenously generated oxidative radicals (**Figs. 2, 3)**, but also altering overall redox balance (including of co-factors) within the cell. Our results therefore link parkin’s own oxidation to concrete redox benefits in cells and organs, thus explaining mechanistically why parkin functions as a ‘multipotent, protective protein’^35^.

Three, parkin’s cellular protection from ROS appeared independent of E3 ligase activity. Its oxidation had previously been widely interpreted as a loss of its function^36,37^, Here, we demonstrated that parkin confers cellular protection and ROS mitigation under oxidizing conditions that would abrogate its E3 ligase activity^9^. We also determined that HMW forms of parkin, which were produced during oxidizing conditions, collapsed when exposed to DTT; this reversal would not occur for parkin species formed by auto-polyubiquitination^38^. Our findings could suggest that E3 ligase function is not essential to parkin’s oxidative stress-mitigating role. However, parkin’s redox-regulating activity may not be mutually exclusive from its E3 ligase function. It is known that low levels of oxidation can activate E3 ligase activity *ex vivo*; moreover, both cellular redox-regulation and E3 ligase function are thiol-based^9,36,39,40^. In support of such a low level of parkin activation, we noted that the protein is oxidized under basal conditions in both murine (**Extended Data Table 1**) and human brain^10^. Furthermore, the reversible nature of some (but not all) of these oxidative modifications suggests that parkin continuously undergoes oxidative events *in vivo*. Therefore, we propose that distinct cellular conditions may require more of parkin’s redox functions, such as during elevated oxidative stress, whereas others may demand more E3 activity, such as during organelle-specific autophagy. Because powerful oxidative stressors are required to recapitulate -in mice-parkin’s aggregation and insolubility (**Fig. 1**), which occurs physiologically during human brain ageing^10^, we speculate that parkin is exposed to significantly higher oxidizing conditions in long-lived humans.

Four, we uncovered a reciprocal relationship between *PRKN* expression and glutathione’s redox state *in vivo*, and generated evidence that parkin plays a direct role in GSSG recycling. The thiol network, to which we propose parkin contributes, is critical in the maintenance of overall redox balance, and thus, to cellular and organ health^41,42^. Here, we expand on the observation made by Itier et al. by comparing a variety of cellular and mouse models to human brain (**Fig. 3**). In parkin’s absence: i) GSH is elevated to increase the total number of free thiols; ii) GSSG levels are diminished; iii) the ratio of GSH:GSSG is increased; and iv) total glutathione levels (GSH + GSSG) are unchanged, except in bigenic mice (**Fig. 3c-f, h; Extended Data Fig. 4a-c, e**). We postulate that parkin’s presence reduced cellular needs for higher levels of free GSH. We first explored *de novo* synthesis of GSH but found no increase in *GCLM* or *GCLC* mRNA levels. As for the rise in total glutathione in bigenic mice, we reasoned that higher GSH levels may still have resulted from *de novo* synthesis but via more effective translation of existing *GCLM* mRNA^28,31^, or elevated enzymatic activity, rather than by gene transcription. By comparing results from the HPLC *vs*. GR-based quantification methods in murine brain, we observed an increase in GSH levels and reciprocal decrease in GSSG levels in *prkn*^*-/-*^ mice when using the latter method, which was therefore attributable to an increase in GR enzyme-related activity (**Fig. 4a, f**)^30,31^. Because we had observed dynamic changes in GSSG levels in cell models following increased parkin expression and oxidative stress (**Fig. 3c-e**), we next explored whether parkin contributed to increased GSH levels by direct thiol-mediated recycling of GSSG. There, we found that parkin directly bound to GSSG in a concentration-dependent manner, which resulted in half of GSSG recycled to GSH and the other half attached to parkin via S-glutathionylation (**Fig. 4b-e**).

The finding of parkin S-glutathionylation (**Fig. 4b, c**), and its reversal by glutaredoxin-1 and -2, further highlights the relevance of parkin’s integration into redox chemistry. The oxidation of proteins via S-glutathionylation represents a major contribution to redox signaling and generally serves as a protective mechanism.^43,44^ We noted with interest that primate-specific cysteine 95 is targeted for oxidation by both S-glutathionylation (**Fig. 4e**) and dopamine radical adduct formation^10^. We question whether this specific residue, which is located in the linker region of human parkin, could have an effect on parkin activation akin to S65 phosphorylation during mitophagy. We anticipate that the many posttranslational modifications of parkin, which we have identified here by LC-MS/MS, will inform structurally oriented and cell biological research in the future. Such investigations, which may focus on the linker region^33^ and downstream mitochondrial health^45^, should be explored in the context of parkin redox chemistry. Due to the transient nature of S-glutathionylation (as well as of S-nitrosylation^39^) as a posttranslational modification and due to the analytical methods used here (**Extended Data Table 1**) and in our analyses of human brain^10^, namely under reducing (and alkylating) conditions, we have not yet validated parkin S-glutathionylation *in vivo*. We hope to continue these investigations in the near future.

Future studies will also explore which region and which cell type in the nervous system^27^ contribute most significantly to the parkin-dependent glutathione redox changes we have observed *in vivo*. The lack of significantly elevated GSH levels in *prkn*^*-/-*^ mouse brain by HPLC suggests that this species can compensate for increased cytosolic ROS via other redox state-regulating mechanisms, such as via glutathione peroxidase (GPx) activity (**Fig. 4f**). High tolerance to ROS may help explain the lack of dopamine cell loss in *prkn*-null mice^18,26,46^. In aged mouse brain, one or more additional stressor could be required, such as demonstrated in the *prkn*^-/-^//*Sod2*^+/-^ mice. A human-specific redox aspect is that nigral neurons experience high levels of ROS production, dysregulated iron homeostasis, and more radical formation from oxidative dopamine metabolism^10,47,48^. Differences in human *vs*. murine parkin oxidative metabolism will thus be further explored.

The strengths of our investigations are three-fold: 1) results in cellular and murine models restaged findings in human tissues^10,19^ (**Figs. 2h, i; 3f, g**); 2) our research integrated *in vitro, ex vivo, in situ* and *in vivo* paradigms as well as exogenous and endogenous ROS sources; and 3) the redox hypothesis of parkin function readily explains its many reported roles in distinct cellular paradigms, *e*.*g*., cancer biology, mitochondrial health, immunity, etc.^49^, because these pathways are (co-)regulated by redox state changes (**Fig. 4f**).

We have also identified shortcomings. First, in future work we will conduct a more extensive assessment of the diversity and quantity of oxidative modifications in brain parkin by LC-MS/MS, for example in our MPTP-exposed *vs*. untreated mouse model, in order to map additional *bona fide* oxidation sites that occur *in vivo*. As mentioned, we have yet to validate S-glutathionylation of parkin *in vivo*, which will require optimization of mass spectrometry techniques. Furthermore, we wish to study a greater number of *PRKN-*genotyped, age-matched, human brain tissue specimens to quantify protein carbonylation and nitrotyrosination, as well as to explore human brain-specific changes in redox state-related transcriptomes from different areas of the brain. Recently, we have begun to characterize glutathione-regulating enzyme levels *in vivo, e*.*g*., glutathione-reductase, to validate our model that its activity is markedly increased in ARPD (**Fig. 4f**).

Our discovery that parkin’s seemingly diverse activities are anchored in traditional redox chemistry (**Fig. 4f**) provides at once a biochemical and unifying explanation for its role in neuroprotection. It may also help answer the two decades-old question: “What function of parkin is essential in conferring the selective protection of dopamine neurons in the adult human brainstem?”^10,50^ Further exploration of parkin’s redox effects should create new opportunities to develop urgently needed therapies for patients with young-onset, currently incurable parkinsonism.

## MATERIALS AND METHODS

### Mouse tissues

Brains and hearts were collected from wild-type C57Bl/6J from Jackson laboratories, *prkn*-null from Dr. Brice’s laboratory ^26^, *Sod2* +/-mice from Jackson laboratories; the bigenic mouse (*prkn*^-/-^//*Sod2*^+/-^) was created by crossing *prkn*-null mice with *Sod2* haploinsufficient mice, and interbreeding heterozygous offspring. The bigenic mouse will be further characterized elsewhere (El Kodsi et al., in preparation). Mouse brains collected were homogenized on ice in a Dounce glass homogenizer by 20 passes in Tris salt buffer with or without the addition of 1% H_2_O_2_, transferred to ultracentrifuge tubes and spun at 55,000 and 4°C for 30 mins to extract the soluble fraction. The resulting pellets were further homogenized in the tris salt buffer with the addition of 2-10% SDS, transferred to ultracentrifuge tubes and spun at 55,000 rpm and 10°C for 30 minutes to extract the insoluble fraction.

### Cell culture, transfection and oxidation

Human embryonic kidney (HEK293) were grown in Dulbecco’s Modified Eagle Medium (DMEM) supplemented with 1 % penicillin/streptomycin and 10 % heat-inactivated fetal bovine serum (FBS) at 37°C with 5 % CO_2_. Four to 15 µg of cDNA coding for N-terminally flag-tagged parkin or empty Flag control vector (pcDNA3.1+) using a 1:1 ratio of cDNA:Lipofectamine 2000, was used for ectopic overexpression. The cDNA and Lipofectamine 2000 were incubated for 20 min at room temperature before being applied directly to the cells for 1 hour at 37°C with 5 % CO_2_ followed by direct addition of fresh growth medium. Cells were incubated another 24 hours before treatment, harvesting and analysis. Control Chinese hamster ovary cells, stably expressing the myc-vector (CHO), or expressing myc-parkin (CHO-parkin) were also used.

All chemicals (H_2_O_2_, CCCP, NEM, IAA, DTT, AT, BSO and NAC) were added directly to cells at ∼75% confluence in growth or OPTI-MEM media. Cells were manually scrapped, spun at a 1000 rpm for 5 minutes, the pellets washed with PBS and then homogenized in a Tris salt buffer, transferred to ultracentrifuge tubes and spun at 55,000 rpm and 4°C for 30 minutes to extract the soluble fraction. The resulting pellets were further homogenized in the Tris salt buffer with the addition of 2-10% SDS, transferred to ultracentrifuge tubes and spun at 55,000 rpm and 10°C for 30 minutes to extract the insoluble fraction. SH-SY5Y cells were generally seeded at a density of 0.5-1 × 106 cells/mL. Once cells reached 70-80 % confluency they were transfected with cDNA coding for C-terminally Flag-tagged Parkin or empty Flag control vector (pcDNA3) by electroporation using the nucleofector method described by Hu and Li, 2015. A total of 2 million cells were resuspended in 100 µL of OPTI-MEM containing cDNA (2 µg) and 1 % polyoxamer 188. The cells were electroporated using the X Unit and pulse code “CA-137” on a Lonza 4D-Nucleofector. Following electroporation, cells were seeded at a concentration of 0.8-1 × 106 cells/mL ^51^.

### Western blot and densitometry

Brain homogenates and cell lysates were run on 4-12 % Bis-Tris SDS-PAGE gels using MES running buffer. Proteins were transferred to PVDF membranes using transfer buffer, and immunoblotted for parkin, DJ-1, MnSOD, aconitase-2, creatine kinase, VDAC, TOM20, nitrotyrosine, and flag-epitopes. Actin and Ponceau S staining were used as loading controls. For densitometry quantification, the signal intensity of parkin from each sample was measured as pixel using Image J Software, and controlled for loading, against actin’s band intensity. Proteins were also stained in gel using Coomassie brilliant blue R-250 dye. The gel was first fixed in 10% acetic acid, 50% methanol, followed by staining for 2-24 hours with Coomassie and finally destained in 7.5% acetic acid, 5% methanol until crisp blue bands appeared.

### MPTP treatment

Eight to 12 months old WT and *prkn*-null mice were injected intraperitoneally with 40mg/kg of saline or MPTP and sacrificed an hour later ^20^. The brains were harvested for ROS measurement, protein analysis by Western blot and immunoprecipitation of parkin and mass spectrometry analysis. For mass spectrometry, the brains harvested were first incubated in IAA prior to homogenization and fractionation as described above. Brain homogenates were then incubated with anti-parkin conjugated to magnetic beads (Dynabeads Coupling Kit; Invitrogen). A magnet was used to capture parkin bound to the beads, and several washes were used to remove unbound proteins. Eluted fractions (IP elute) along with controls (input, unbound, wash and recombinant parkin protein standards) were run on SDS/PAGE and blotted with anti-parkin. A sister gel was stained with Coomassie as described above and gel slices corresponding to band sizes 50-75 kDa were excised and analyzed by LC-MS/MS, as described in detail by Tokarew et al., 2020.

### Recombinant, tag-less protein expression in pET-SUMO vector

Wild-type and truncated (residues 321-465) human parkin were expressed as 6His-Smt3 fusion proteins in *Escherichia coli* BL21 (DE3) Codon-Plus RIL competent cells (C2527, New England Biolabs) as previous described^22,33,52^. DJ-1 and SNCA coding sequences were cloned from a pcDNA3.1 vector into the pET-SUMO vector using PCR and restriction enzymes. All proteins were overexpressed in *E. coli* BL21 Codon-Plus competent cells (C2527, New England Biolabs) and grown at 37 °C in 2 % Luria Broth containing 30 mg/L kanamycin until OD600 reached 0.6, at which point the temperature was reduced to 16°C. Parkin protein-expressing cultures were also supplemented with 0.5 mM ZnCl_2_. Once OD600 reached 0.8, protein expression was induced with isopropyl β-D-1-thiogalactopyranoside, except ulp1 protease, which was induced once OD600 had reached 1.2. The concentration of IPTG used for each construct is as follows: 25 µM for wild-type and point mutants of parkin, and 0.75 mM for truncated parkin, DJ-1, α-synuclein, SAG, and ulp1 protease. Cultures were left to express protein for 16-20 h. Cells were then harvested, centrifuged, lysed and collected on Ni-NTA agarose beads in elution columns.

### Recombinant maltose-binding protein-tagged protein expression in pMAL-2T vector

Wild-type and truncated (residues 327-465) human parkin were expressed in the pMAL-2T vector (a gift from Dr. Keiji Tanaka), as previously described^53^. Parkin produced in this vector contained an N-terminal maltose-binding protein (MBP) and thrombin cleavage site (LVPRGS). All proteins were overexpressed in *E. coli* BL21 Codon-Plus competent cells (C2527, New England Biolabs) and grown at 37 °C in 2 % Luria Broth containing 0.2 % glucose and 100 mg/L ampicillin until OD600 reached 0.3-0.37, at which point protein expression was induced with addition of 0.4 mM isopropyl β-D-1-thiogalactopyranoside. Cultures were left to express protein at 37 °C until OD600 reached 0.9-1.0. Harvested protein isolates were purified using amylose resin in buffers containing 100 µM zinc sulfate and 10 mM maltose.

### ROS (H_2_O_2_) measurements of recombinant proteins, tissue and cell lysates

Amplex^®^ Red hydrogen peroxide/peroxidase assay kit (Invitrogen A22188) was used to monitor endogenous levels of H_2_O_2_ in tissues and cells, and residual levels of H_2_O_2_ after incubation with recombinant parkin (WT, or pre-incubated with increasing concentrations of H_2_O_2_, NEM, or EDTA), DJ-1, SNCA, BSA, RNF43 (from BioLegend), HOIP (from Boston Biochem), GSH, catalase, NEM and EDTA for 30 minutes. Pre-weighed cortex pieces from human brains (or pelleted cells) were homogenized on ice in the 1x reaction buffer provided, using a Dounce homogenizer (3 times volume to weight ratio). Homogenates were diluted in the same 1x reaction buffer (10x and 5x). A serial dilution of the H_2_O_2_ standard provided was prepared (20, 10, 2 and 0 µM). 50 µL of standards and samples were plated in a 96 well black plate with clear flat bottom. The reaction was started by the addition of 50µL working solution which consist of 1x reaction buffer, Amplex^®^ red and horseradish peroxidase. The plate was incubated at room temperature for 30 minutes protected from light. A microplate reader was used to measure either fluorescence with excitation at 560 nm and emission at 590 nm, or absorbance at 560 nm. The obtained H_2_O_2_ levels (µM) were normalized to the tissue weight (g) or protein concentration (µg/µL). The same aasay was also used to measure parkin and glutathione’s peroxidase activity compared to horseradish peroxidase (HRP).

### ROS (H_2_O_2_) measurements of intact cells

HEK293 cells were transfected with flag-parkin or control vector (pcDNA) as described above. After 24 h the cells were lifted using trypsin and re-seeded in a 12-well dish at a density of 0.3 × 106 cells/mL. After 48 h the cells were treated with 0 mM or 2 mM H_2_O_2_ in OPTI-MEM medium at 37°C and 5 % CO_2_. After 1 h the cells were washed with OPTI-MEM and incubated with 20 µM of dichlorofluorescin diacetate (DCFH-DA, D6883, Sigma) for 30 min at 37°C and 5 % CO_2_. Cells were collected using a cell lifter and treated with ethidium-1 dead stain (E1169, Invitrogen) for 15 min at room temperature. Samples were analyzed using a BD Fortessa flow cytometer set to measure the ROS-sensitive probe (DCFH-DA, ex. 488 nm and em. 527 nm) and viability stain (ethidium-1, ex, 528 nm and em. 617 nm). The results were reported as the average mean fluorescence intensity (MFI) of ROS in live cells. Each separate transfection was considered one biological replicate.

### Cell cytotoxicity assay

Vybrant ™ cytotoxicity assay kit (Molecular Probes V-23111) was used to monitor cell death through the release of the cytosolic enzyme glucose 6-phosphate dehydrogenase (G6yPD) from damaged cells into the surrounding medium. 50µl of media alone (no cells), media from control and stressed CHO-parkin and control cells and cell lysates were added to a 96-well microplate. 50 µl of reaction mixture, containing reaction buffer, reaction mixture and resazurin, was added to all wells, and the mircroplate was incubated at 37°C for 30 mins. A microplate reader was used to measure either fluorescence with excitation at 560 nm and emission at 590 nm. A rise in fluoresence indicates a rise in G6PD levels i.e. a rise in cell death.

### Mitochondria isolation

Fresh tissue was cut with scissors into manageable pieces, rinsed in cold PBS, then homogenized using either a Dounce homogenizer or Warning blender in the presence of twice the tissue volume of buffer A (20mM Hepes pH 7.4, 220mM mannitol, 68mM sucrose, 80mM KCl, 0,5mM EGTA, 2mM Mg(Ac)_2_, 1mM DTT, 1X protease inhibitor (Roche)). The sample was centrifuged at 4070g in a tabletop centrifuge for 20 minutes at 4°C. The supernatant was collected and spun again as above. The resulting supernatant was spun at 10 000g for 20 minutes at 4°C and the pellet was then washed in the above buffer and spun again at 10 000g for 20 minutes. The resulting mitochondrial pellet was carefully resuspended in buffer B (20mM Hepes pH 7.4, 220mM mannitol, 68mM sucrose, 80mM KCl, 0,5mM EGTA, 2mM Mg(Ac)_2_, 10% glycerol), aliquoted, and snap frozen.

### Aconitase assay

The Aconitase Enzyme Activity Microplate Assay Kit (MitoSciences) was used to measure aconitase activity in mitochondria isolated from wild type and parkin knock-out mouse brains as per manufacturer’s instructions. Two brains each from 12 months old mice were pooled to provide the mitochondria samples and normalized for total protein concentration. These were treated with 0 or 4 μM H_2_O_2_ just prior to assay. The catalytic conversion of isocitrate to cis-aconitate by aconitase was measured by quantifying the amount of cis-aconitate in the reaction by reading the samples at 240nm. Rates in μM/min were determined from 3 independent experiments performed in triplicate.

### Creatine kinase assay

The EnzyChrom Creatine Kinase Assay Kit (BioAssay Systems) was applied to measure creatine kinase activity in mitochondria purified from wild type and parkin knock-out mouse brains following the manufacturer’s instructions. Two brains were pooled each from 12 months old mice to provide the mitochondrial samples and normalized for total protein concentration. Mitochondria were incubated with 0 or 0.5mM H_2_O_2_ at room temperature for 20 minutes prior to assay. The creatine kinase-dependent catalytic conversion of creatine phosphate and ADP to creatine and ATP, was quantified indirectly by measuring NADPH at 340nm. The ATP produced by the reaction phosphorylates glucose to glucose-6-phosphate by hexokinase, which is then oxidized by NADP in the presence of glucose-6-phosphate dehydrogenase, yielding NADPH. Rates in μM/min were calculated for 3 independent experiments done in triplicate.

### GSH and GSSG quantification - HPLC

Human and mouse brain pieces, and CHO cell pellets were homogenized in buffer containing 125 mM sucrose, 5 mM TRIS, 1.5 mM EDTA, 0.5%TFA and 0.5%MPA in mobile phase. Then samples were spun at 14000xg at 4 °C for 20 mins. Supernatants were collected and analyzed using an Agilent HPLC system equipped with a Pursuit C_18_ column (150 ×4.6 mm, 5 µm; Agilent Technologies) operating at a flow rate of 1 ml/min or. The mobile phase consisted of 0.09% trifluoroacetic acid diluted in ddH2O and mixed with HPLC-grade methanol in a 90:10 ratio. Standard solutions were used to estimate the retention times for GSH and GSSG. Using Agilent Chemstation software, absolute amounts of GSH and GSSG were acquired by integrating the area under the corresponding peaks, and values were calculated from standard curves

### GSH concentration determination by monochlorobimane assay

Stock solutions of assay dye (monochlorobimane (MCB), 22 mM) and glutathione-S-transferase (50 units/mL) were prepared in PBS and stored protected from light at −20°C. The working solution was prepared using 12.8 µL of stock mcB and 80 µL of stock glutathione-S-transferase in 4 mL PBS and stored on ice. Samples were prepared as follows: cells were lifted mechanically using cell-lifters, washed twice and re-suspended in ice-cold PBS, mixed by vortex and incubated on ice for 30 min. Following two freeze thaw cycles using solid CO_2_, the samples were sonicated 1 min on wet ice (S220 Ultra-sonicator from Covaris) and spun at 3000 x g, 4°C, for 5 min. Total protein concentration of supernatants was determined using Bradford assay. Samples and glutathione (GSH) standards (0-13 µM) were plated in 25 µL aliquots in a 96-well plate with clear bottom and black sides. 25 µL of working solution was added to all experimental wells and protected from light for 15 min at room temperature. Fluorescence (ex 380 nm, em 461 nm) was measured using a Synergy H1Multi-Mode Plate Reader (Bio Tek). The amount of GSH detected in each sample was calculated using the regression curve obtained from the glutathione standards.

### Tietze’s enzymatic recycling determination of GSH and GSSG

The enzymatic recycling method described by Rahman et al.^54^ was used to determine reduced glutathione (GSH) and oxidized glutathione (GSSG) levels in mouse brain lysates. Hemi-brains of wild type (n=3) and parkin KO (n=3) mice, at 13 and 11 months of age respectively, were collected, weighed and homogenized in 3X volume/weight of KPEX (0.1 M potassium phosphate, 5 mM EDTA, 0.1 % Triton X-100, 0.6 % sulfosalicylic acid, pH 7.5) using a glass Dounce homogenizer (50 passes). Samples were spun at 8000 x g, 4°C, for 5 min and the supernatant protein concentration was determined using Bradford assay. To determine the total glutathione (GSH + GSSG) concentration, the following stock solutions were freshly prepared in KPE (0.1 M potassium phosphate, 5 mM EDTA, pH 7.5): 5,5’-dithio-bis-[2-nitrobenzoic acid] (DNTB) at 0.6 mg/mL, nicotinamide adenine dinucleotide phosphate (NADPH) at 0.6 mg/mL and glutathione reductase at 3 units/mL. GSH standards were prepared in KPE at concentrations of 0-26 nM/mL. 20 µL of diluted sample or GSH standard was added per well and 120 µL of a 1:1 mixture of the DNTB and glutathione reductase stocks solutions was added to each assayed well. After 30 sec incubation, 60 µL of the NADPH was added and absorbance was immediately measured at 412 nm in 30 sec intervals for a total of 2 min. To determine the concentration of oxidized glutathione (GSSG), the samples were first diluted (1 in 4) in KPE and treated with 0.2 % 2-vinylpyridine for 1 h at room temperature. Excess vinyl-pyridine was quenched with 1 % triethanolamine and GSSG was measured using the same method as total glutathione except GSH standards were replaced with GSSG standards (0-26.24 nM/mL) that were treated with vinyl-pyridine and triethanolamine. The absolute values of total glutathione (GSH + GSSG) and oxidized glutathione (GSSG) per sample were calculated using the linear regression obtained from the change in absorbance/min plotted against the GSH or GSSG standard concentrations, respectively, and dividing by the total protein concentration. Absolute values for GSH were determined using the following equation: GSH= [GSH + GSSG] – 2[GSSG] ^54^.

### Parkin-catalyzed redox state recycling

Parkin protein buffer exchange to T200 protein buffer (50 mM Tris, 200 mM NaCl, pH 7.5) was first performed using repeat centrifugations (8 times 4000 x g at 4°C for 10 min) in Amicon Ultra 10 kDa MWCO filters. Protein concentration was adjusted to 10 µM using T200. Both reduced (GSH) and oxidized (GSSG) glutathione stocks were prepared in phosphate buffered saline at concentrations of 1 mg/mL (3250 µM) and 2.01 mg/mL (6560 µM) respectively. Glutathione standards of 0, 2.5, 5, 10 µM and 100 µM of both GSH and GSSG were prepared and combined in the following ratios to a final volume of 90 µL at: 10 µM GSH: 0 µM GSSG, 9 µM GSH: 1 µM GSSG, 8 µM GSH: 2 µM GSSG, 6 µM GSH: 4 µM GSSG, 4 µM GSH: 6 µM GSSG, 2 µM GSH: 8 µM GSSG, 1 µM GSH: 9 µM GSSG, and 0 µM GSH: 10 µM GSSG. R-parkin (1 µL of a 10 µM solution) was added to the prepared mixtures and allowed to incubate at room temperature for 15 min. Samples were analyzed for GSH concentration using the monochlorobimane assay described above.

### Glutathionylation assay

Glutathionylation of tagged and untagged parkin proteins was performed, as described previously ^55^. MBP-tagged parkin proteins were eluted from columns with excess maltose. Concentrated eluates were supplemented with 0.1% DMSO (10 µl DMSO in 10 ml PBS), and excess DTT and maltose were removed by several cycles of centrifugation with 30 kDa cut-off filters. Proteins/peptides (14 µM) were incubated with 3 mM GSH for 1 h and then with 5 mM GSSG for 2 h at room temperature. Trypsin digestion was performed (Peptide: Trypsin = 20: 1) overnight at 4 °C. The Trypsin-digested fragments were run through MALDI analysis. To assay S-glutathionylation, the eosin-labeled GSSG (Di-E-GSSG) was used to glutathionylate proteins, as described previously ^32^. Di-E-GSSG has quenched fluorescence in the disulphide form, which increases ∼20-fold upon reduction of its disulphide bond and formation of E-GSH. Blackened 96-well-plates were used in a PerkinElmer Victor3 multilabel counter containing a final well volume of 200 μl in 0.1 M potassium phosphate buffer (pH 7.5), 1 mM EDTA. The reaction was started by addition of 20 μM Di-E-GSSG to parkin proteins, followed by recording the fluorescence emission at 545 nm after excitation at 520 nm. Controls with no peptide added were used as fluorescent background. To confirm S-glutathionylation, reaction products were incubated with Di-E-GSSG. Aliquots of S-glutathionylated proteins were further treated with 10 mM DTT or with the complete GSH-Grx system. All samples were run on a non-reducing SDS-PAGE 4–12% acrylamide. The gel was exposed to UV transilluminator to visualize eosin-tagged glutathionylated protein. The same gels were later stained with Coomassie Blue staining. Di-Eosin-GSSG was purchased form IMCO, Sweden. Human Grx1, and Grx2 were prepared as described previously ^32^. Rat recombinant TrxR was a kind gift from Prof. Elias Arner.

### Mass spectrometry analysis of glutathionylated parkin

The protein was treated with proteolytic digestion, using trypsin. The resulting peptides were separated using one dimension of liquid chromatography. The LC eluent was interfaced to a mass spectrometer using electrospray ionization and the peptides were analyzed by MS.

LC−MS/MS analyses were performed using an Easy-nLC chromatography system directly coupled online to a Thermo Scientific Q Exactive hybrid quadrupole-Orbitrap mass spectrometer with a Thermo Scientific(tm) Nanospray Flex(tm) ion source. The sample was injected from a cooled autosampler onto a 10 cm long fused silica tip column (SilicaTips, New Objective, USA) packed in-house with 1.9 μm C18-AQ ReproSil-Pur (Dr. Maisch, Germany). The chromatographic separation was achieved using an acetonitrile (ACN)/ water solvent system containing 0,1% FA and a gradient of 60 min from 5 to 35% of ACN. The flow rate during the gradient was 300 nL/ min.

MS/MS data were extracted and searched against in-house Mascot Server, (Revision 2.5.0), search engine that uses mass spectrometry data to identify and characterize proteins from sequence databases. The following parameters were used: trypsin digestion with a maximum of two missed cleavages; Carbamidomethyl (C), Oxidation (M), Deamidated (NQ) and Glutathione (G) as variable modifications; and a precursor mass tolerance of 10 ppm and a fragment mass tolerance of 0.02 Da. The identified protein was filtered using 1% false discovery rate (FDR) and at least two peptides per protein as limiting parameters.

### Protein carbonyl assay

Protein carbonyl colorimetric assay kit (Cayman chemical 10005020) was used to assay the carbonyl content in human and mouse brains or hearts. Pre-weighed tissues were rinsed in PBS and then homogenized in 1 mL cold PBS at pH 6.7 supplemented with 1 mM EDTA, using a Dounce homogenizer on ice. Homogenates were centrifuged at 10,000xg for 15 minutes at 4 °C. 200 µL of the supernatant was added to a tube with 800 µL DNPH (sample tube) and 200 µL of the supernatant was added to a tube with 800 µL 2.5 M HCL (control tube), both tubes were incubated in the dark for 1 hour with occasional vortex. 1 mL 20 % TCA followed by 1 mL 10% TCA solutions were added to centrifuged (10,000xg 10 minutes at 4 °C) pellet after discarding the supernatant. The resulting pellet was resuspended in1 mL of 1:1 ethanol:ethyl acetate mixture and centrifuged 3 times to extract protein pellets. The final pellets were suspended in 500 µL guanidine hydrochloride and centrifuged. A total of 220 µL per sample and control supernatants were added to two wells of a 96-well plate, and the absorbance was measure at 360 nm. The corrected absorbance (sample – control) was used in the following equation to obtain the protein carbonyl concentration: Protein Carbonyl (nmol/mL) = [(CA)/(0.011 μM^-1^)](500 µL/200 µL). Total protein concentration from the sample tissues were measured to obtain the carbonyl content: protein carbonyl/total protein concentration.

### MnSOD activity assay

Superoxide dismutase assay kit (Cayman chemical 706002) was used to assay MnSOD activity in mouse brains. Pre-weighed perfused mouse brain pieces were homogenized in 5 mL cold 20 mM HEPES buffer, pH 7.2, supplemented with EGTA, mannitol and sucrose, with a Dounce homogenizer on ice. The homogenates were centrifuged at 1,500xg 5 minutes at 4 °C. The SOD standards were prepared by adding 200 µL of the radical detector and 10 µL of the provided standards, in duplicates in a 96-well plate. The same was repeated for the samples. The reaction was initiated by adding 20 µL of xanthine oxidase to all the wells. Background absorbance was assayed by adding 20 µL xanthine oxidase to sample buffer (optional). The plate was incubated on a shaker for 30 minutes at room temperature. The absorbance was measure at 450 nm. The linearized SOD standard curve was plotted and used to calculate the MnSOD activity (U/mL) from the averaged sample absorbances.

### Total RNA isolation, cDNA synthesis and PCR amplification

Pre-weighed cortex pieces from mouse brains were homogenized in QIAzol (Qiagen 79306), at 1ml volume per 100mg of tissue, and incubated at room temperature for 5 minutes. 0.2mL of chloroform (per 1mL QIAzol) was added and the homogenates were shaken vigorously for 15 seconds, followed by a 2-3 minutes incubation at room temperature, the tubes were centrifuged at 12,000xg for 15 minutes at 4°C. The upper clear aqueous layer was transferred to a new tube and 1 volume of 70% ethanol was added, and mixed by vortexing. The solution was then added to an RNAeasy Mini spin column (Qiagen 74104) placed in a 2mL collection tube and centrifuged for 15 sec at 8000xg at room temperature. The flow-through was discarded and 700 µL Buffer RW1 was added to the spin column and spun for 15 sec at 8,000xg. The same step was repeated with 500 µL Buffer RPE, one spin for 15 sec and a second spin for 2 min. An optional spin in a new collection tube at full speed for 1 min to remove excess buffer was be added. The RNAeasy Mini spin column was placed in a new collection tube, 50 µL RNase-free water was added directly to the membrane and centrifuged for 1 minute at 8000xg. A NanoDrop machine was used to measure the amount of total RNA obtained from the cortices. Turbo DNA-free™ (Life Technologies 1412059) was used to remove trace to moderate amounts of contaminating DNA. SuperScrpit™ IV First-Strand Synthesis System (Invitrogen 18091050) was used for cDNA synthesis reaction. iTaq™ Universal SYBR® Green Supermix (BIO-RAD 172-5121) and select primer sets were used for PCR amplification of the newly synthesized cDNA templates, and controls, and analyzed by agarose gel electrophoresis and ethidium bromide staining. The following primers were used ^56-58^:

DJ-1, F: ATCTGAGTCGCCTATGGTGAAG; R: ACCTACTTCGTGAGCCAACAG

GCLC, F: ATGTGGACACCCGATGCAGTATT; R: TGTCTTGCTTGTAGTCAGGATGGTTT

GCLM, F: GCCACCAGATTTGACTGCCTTT; R: CAGGGATGCTTTCTTGAAGAGCTT

Actin, F: CTTCCTCCCTGGAGAAGAGC; R: AAGGAAGGCTGGAAAAGAGC

### Statistical analyses

All statistical analyses were performed using GraphPad Prism version 8 (GraphPad Software, San Diego, CA, USA, www.graphpad.com). Differences between two groups were assessed using an unpaired t-test. Differences among 3 or more groups were assessed using a 1way or 2way ANOVA followed by Tukey’s post hoc corrections to identify statistical significance. Subsequent post hoc tests are depicted graphically and show significance between treatments. For all statistical analysis a cut-off for significance was set at 0.05. Data is displayed with p values represented as *p < 0.05, **p < 0.01, ***p < 0.001, and ****p < 0.0001.

## ACKNOWLEDGMENTS

We are grateful for the commitment of patients and their family members to participate in autopsy studies. We thank Drs. A. Brice and E. Fon for sharing *prkn*-null mice, Drs. N. Matsuda and K. Tanaka for MBP-parkin cDNA-encoding plasmids; Drs. R. Tam, S. Bennett and D. Pratt for helpful discussions, and members of the Schlossmacher lab for their critical comments and suggestions.

## Author contributions

*Study design:* D.N.E.K., J.M.T., J.J.T., M.G.S.; *Writing and figure preparation:* D.N.E.K., J.M.T., N.A.L., J.J.T. and M.G.S. prepared the initial draft of the manuscript and figures. All authors reviewed and / or edited the manuscript and approved of the submitted version. *Experiments*: D.N.E.K., J.M.T., N.A.L., R.S., A.C.N., H.B., C.P., B.S., D.S.I., L.Z., and L.P. performed experiments; R.S., C.P, J.A.C, M.T, N.H., R.R.R. and L.P provided data, tissue specimens and critical comments. *Analysis:* D.N.E.K. J.M.T, J.L., J.J.T., and M.G.S. performed data analyses. *Study supervision*: J.J.T. and M.G.S. *Overall responsibility:* M.G.S. This work is dedicated to the memories of Mr. Bruce Hayter (1962-2019), a tireless advocate for persons with young-onset parkinsonism, and our colleague, Dr. Arne Holmgren (1940-2020), a pioneer in redox biology.

## Additional Information

**Data and materials availability**: Original data associated with this study are available in the main text and extended data figures and tables; additional data will be made available upon request.

**Supplementary Information** is available for this manuscript.

**Extended Data Table 1:** Parkin cysteines are oxidized in adult mouse brain. Table summarizing mass spectrometry results highlighting the ratios of the number of IAA-tagged and free cysteines divided by the total number of identified cysteines in wild-type parkin enriched from the brains of mice treated with MPTP (2 independent runs, n=3 brains each) compared to saline controls (2 independent runs, n=3 brains each) under reducing and alkylating conditions (see Methods for details). The means of both runs in each treatment are shown with the ratio (in percent) of IAA-tagged cysteines to total identified cysteines highlighted. Select, oxidative stress-induced, posttranslational modifications found at murine parkin cysteines are highlighted in the table (bottom), *e*.*g*.: cysteine to dehydroalanine; cysteine dioxidation (=sulfinic acid); and cysteine trioxidation (=sulfonic acid). Phosphorylation and methionine oxidation events (table only) are also listed. Graphic summary of murine parkin domains with selected residues for identified PTMs. OX, oxidized cysteines (sulfinic or sulfonic acid); DHA, dehydroalanine; P, phosphorylation.

| Run | 1 |  | 2 |  | Mean |  |
| --- | --- | --- | --- | --- | --- | --- |
| Treatment | Saline (n=1) | MPTP (n=1) | Saline (n=2) | MPTP (n=2) | Saline | MPTP |
| Number of unique parkin peptides identified | 11 | 18 | 11 | 27 | 11.0 | 22.5 |
| Number of amino acids identified | 106 | 154 | 123 | 323 | 114.5 | 238.5 |
| % coverage | 23 | 33 | 27 | 70 | 25.0 | 51.5 |
| Number of cysteines identified | 6 | 10 | 7 | 26 | 6.5 | 18.0 |
| % of total cysteines | 17.6 | 29.4 | 20.6 | 76.5 | 19.1 | 52.9 |
| Number of IAA-modified cysteines identified | 6 | 8 | 7 | 22 | 6.5 | 15.0 |
| Ratio IAA-cysteines/total cysteines identified | 100.0 | 80.0 | 100.0 | 84.6 | 100.0 | 82.3 |
| Cysteine -> Dehydroalanine | C195 | C195 | C195, C240, C252 | C195, C237, C240, C252, C376 |  |  |
| Sulfonic acid (Cys trioxidation) | C252 | C252 | C252, C417, C420 | C211, C252 |  |  |
| Sulfinic acid (Cys dioxidation) | n/d | n/d | n/d | C252 |  |  |
| Phosphorylation | n/d | n/d | n/d | S77 |  |  |
| Methionine oxidation | M191, M457 | M191, M457 | M191, M457 | M191, M326, M457 |  |  |

**Extended Data Figure 1:**
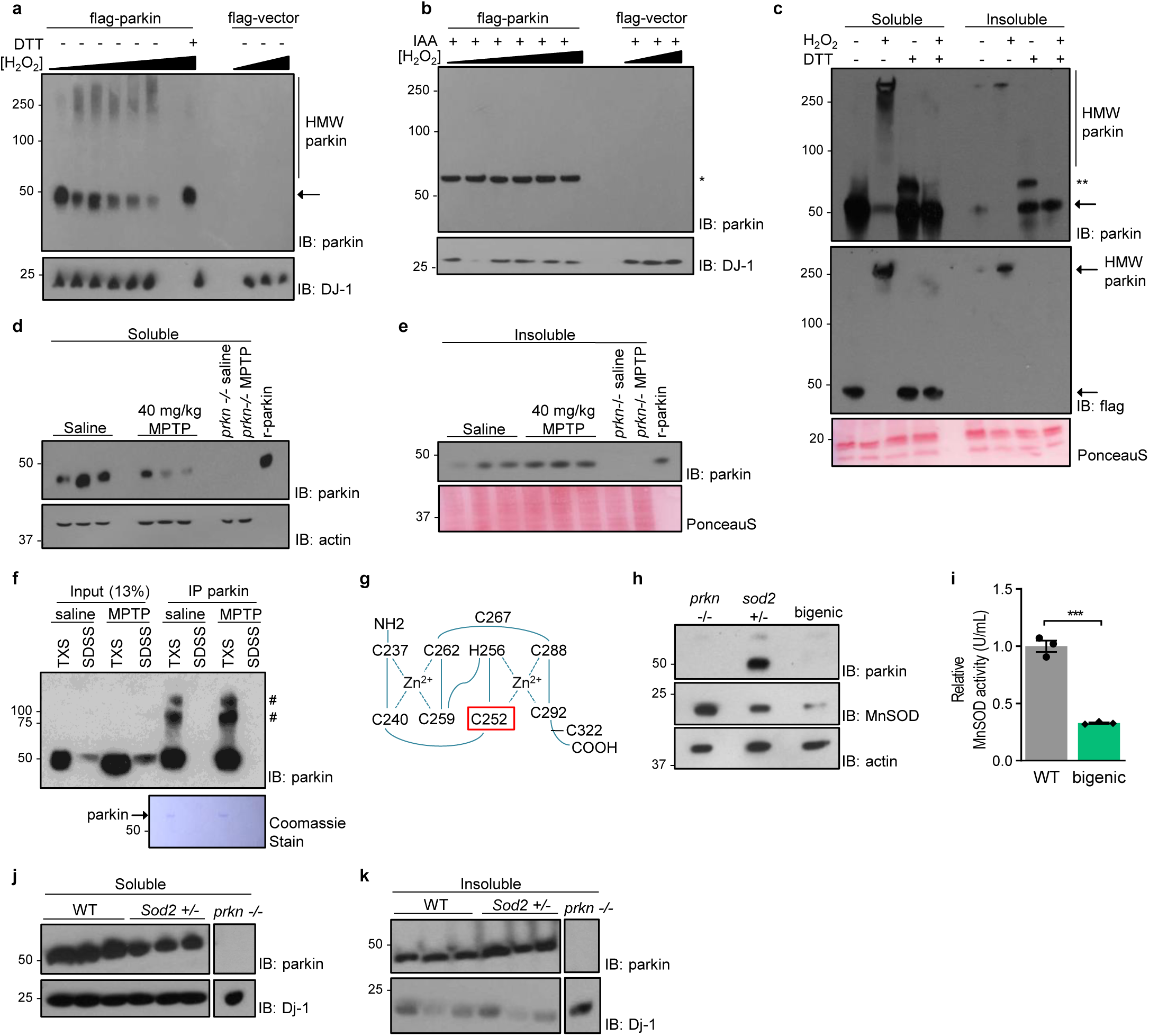
Mitochondrial and cytosolic stressors lead to oxidation of cellular parkin promoting its insolubility. Representative Western blots of parkin and DJ-1 in HEK293-parkin or control cells incubated with increasing concentrations of H_2_O_2_ (2μM to 20mM) **a**, with the addition of dithiothreitol (DTT) (100mM, 30 mins; lane 8) or **b**, pre-incubating with iodoacetamide (IAA) (10mM, 15 mins). **c**, Representative Western blots of parkin and flag in cell lysates from HEK293-parkin or controls cells under oxidizing and/or reducing conditions. (←) Monomeric parkin, (*) Alkylated-parkin and (**) DTT-modified-parkin. Representative Western blots of parkin distribution in **d**, soluble, and **e**, insoluble fractions of whole or hemi brain homogenates from mice treated with saline or MPTP. **f**, Representative Western blots of immunoprecipitated parkin from mouse brains treated with saline or MPTP. (#) IgG related bands. **g**, Schematic representation of parkin’s RING1 domain with the position of an oxidized cysteine residue highlighted (C252). **h**, Representative Western blots of parkin and MnSOD in brains of 2-4 mths-old *prkn*^-/-^, *Sod2*^+/-^ and bi-genic mice. **i**, MnSOD activity in brains of 2-4 mths-old WT and bi-genic mice. Representative Western blots of parkin and Dj-1 distribution in **j**, soluble, and **k**, insoluble fractions of brains from WT and *Sod2*^+/-^ mice. A Student t-test was used for statistical analysis (*** = < 0.001).

**Extended Data Figure 2:**
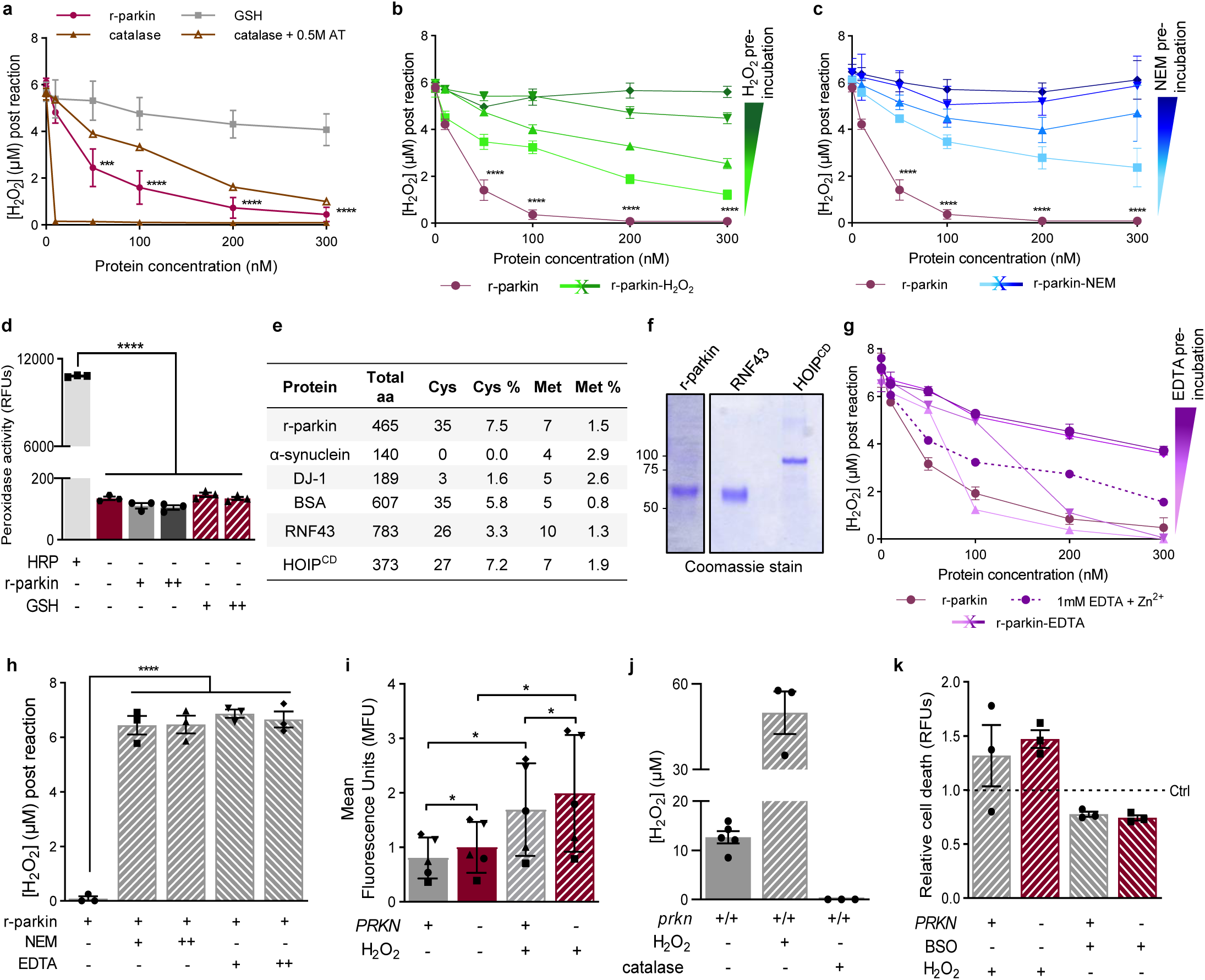
Parkin cysteines reduce hydrogen peroxide to lower oxidative stress. Parkin’s reducing capacity **a**, compared to glutathione and catalase, at equal molar concentrations (AT was used to block catalase activity), **b**, post oxidation with increasing concentrations of H_2_O_2_ (2μM to 2mM), and **c**, after incubation with increasing concentrations of NEM (100μM to 1mM). **d**, Parkin’s and glutathione’s peroxidase activity in comparison to horseradish peroxidase. **e**, Table summarizing the cysteine and methionine content of parkin, DJ-1, α-synuclein, BSA, RNF43 and HOIP^cd^. **f**, r-parkin, RNF43 and HOIP^cd^ proteins visualized by Coomassie stain. **g**, Parkin’s reducing capacity post incubation with increasing concentrations of ethylenediaminetetraacetic acid (EDTA) (100μM to 1mM). **h**, H_2_O_2_ reducing capacity for NEM and EDTA. **i**, H_2_O_2_ levels in live HEK293-parkin or control cells treated with 0 or 2 mM H_2_O_2_ (flow cytometry). Paired data are represented by a distinct symbol. **j**, AmplexRed assay sensitivity demonstrated by the addition of H_2_O_2_ or catalase to mouse brain homogenates. **k**, Cell cytotoxicity in CHO-parkin or control cells with the addition of 2mM H_2_O_2_ or 20mM BSO alone. A 1way, or 2way, unpaired or paired, ANOVA with Bonferroni post hoc analysis were used for statistical analysis (*= <0.05; ** = < 0.01; ***= <0.001; and **** = <0.0001).

**Extended Data Figure 3:**
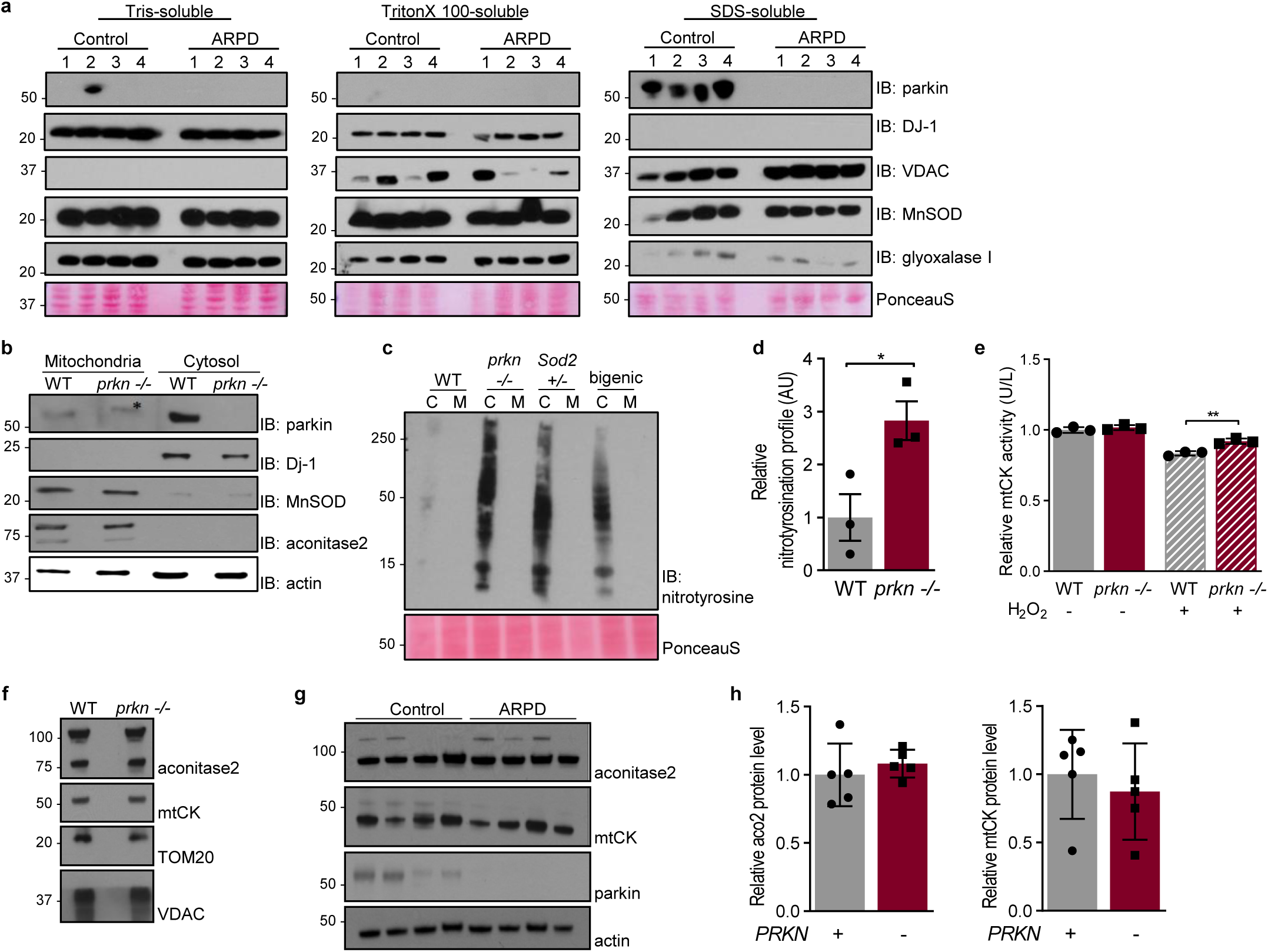
*PRKN* expression alters glutathione metabolism in cells and human brain. **a**, Representative Western blots of parkin, DJ-1, VDAC, MnSOD, and glyoxalase-1 in Tris-soluble, TritonX 100-soluble and SDS-soluble fractions of aged-matched control and human ARPD (*PRKN* mutant) cortices. **b**, Representative Western blots of parkin, DJ-1, MnSOD, and aconitase-2 in mitochondria-enriched vs. cytosol fractions from WT and *prkn*^-/-^ mouse brains. (*) Non-specific band. **c**, Representative Western blots of nitrotyrosination in heart homogenates from 6 mths-old WT, *prkn*^-/-^, *Sod2*^+/-^ and bi-genic mice. **d**, Quantification of nitrotyrosination signals in WT and *prkn*^-/-^ mouse hearts. **e**, Creatine kinase (mtCK) enzymatic activity in mitochondria isolated from 12 mths-old WT and *prkn*^-/-^ mouse brains. Representative Western blots of aconitase-2, mtCK, TOM20, VDAC and parkin in mitochondrial extracts from **f**, mouse brains, and **g**, human brains. **h**, Quantification of aconitase-2 and mtCK signal distribution in mammalian brains (mouse and human). A Student t-test or 1way ANOVA was used for statistical analysis (* = < 0.05; and ** = < 0.01).

**Extended Data Figure 4:**
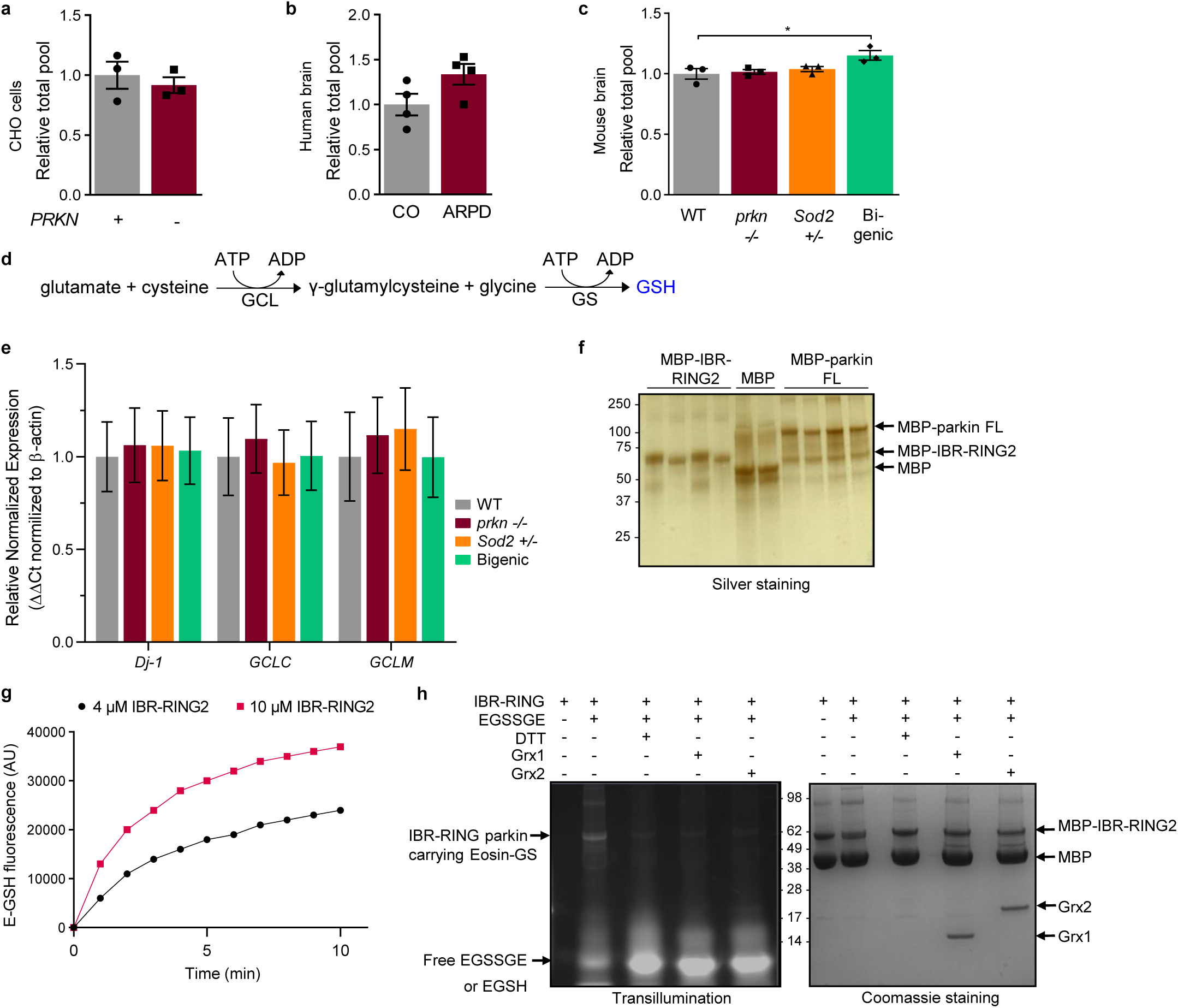
Parkin contributes to glutathione recycling independent of *de novo* synthesis. The total GSH + GSSG pool in **a**, CHO cells, **b**, human cortices, and **c**, brain homogenates of 6 mths-old WT, *prkn*^-/-^, S*od2*^+/-^, and bi-genic mice (HPLC method). A 1way ANOVA was used for statistical analysis (*= <0.05). **d**, Graphic depiction of GSH synthesis from glutamate and cysteine, with glutamate cysteine ligase (GCL) as the rate-limiting enzyme. **e**, *Dj-1*, glutamate cysteine ligase catalytic subunit (*GCLC)* and glutamate cysteine ligase modifier subunit (*GCLM)* gene expression in brains of 6 mths-old WT, *prkn*^-/-^, *Sod2*^+/-^ and bi-genic mice. **f**, Silver stain of MBP-tag (∼45-50 kDa), MBP-IBR-RING2 (parkin, aa 327-465, ∼60-65 kDa), and MPB-parkin (∼90-95 kDa). **g**, E-GSH release post treatment of increasing concentrations of MBP-IBR-RING2 protein with 20μM Di-E-GSSG. **h**, Representative UV transillumination and Coomassie Blue stain gels of S-glutathionylated MBP-IBR-RING2 protein. Results are representative of three independent experiments. Grx1, glutaredoxin-1; Grx2, glutaredoxin-2; EGSSGE, Di-E-GSSG (eosin-labelled oxidized glutathione).

